# Soil microbial and plant responses to increasing antibiotic concentration: a case study of five antibiotics

**DOI:** 10.1101/2025.08.06.668893

**Authors:** Sarah van den Broek, Inna Nybom, Rafaela Feola Conz, Yifei Sun, Thomas D. Bucheli, Sebastian Doetterl, Martin Hartmann, Gina Garland

## Abstract

Antibiotic contamination from biogenic waste in agricultural soils poses a significant threat to soil health and crop productivity. We investigated the effect of antibiotics on the soil microbial community, antibiotic resistance genes (ARGs) and mobile genetic elements (MGEs) and plant productivity in a six week greenhouse trial. Here, *Spinacia oleracea* (spinach) and *Raphanus sativus* (radish) were grown from seed and a mix of five antibiotics, namely sulfamethoxazole, trimethoprim, enrofloxacin, clarithromycin and chlortetracycline, were added to the soil at concentrations 0, 0.1, 1 and 10 mg kg^−1^ soil dry weight (c0, c0.1, c1 and c10, respectively). Overall, we found that the antibiotic treatments significantly impacted prokaryotic α-diversity and prokaryotic and fungal β-diversity. Human and plant pathogen abundance did not increase under antibiotic exposure, but there was a significant reduction of plant growth-promoting bacteria. Moreover, the c10 treatment significantly increased the abundance of MGE *intI1* indicative of horizontal gene transfer and ARG *sul1* antibiotic resistance and significantly lowered radish biomass and nitrogen uptake, while spinach biomass and nitrogen uptake were unaffected. In summary, our study showed that antibiotic exposure significantly changed prokaryotic community diversity and taxonomy, while fungi remained largely unaffected. The reduction of plant growth-promoting bacteria may have a significant impact on soil nutrient cycling and crop productivity, but more research is needed to understand the long-term impact of these co-applied antibiotics on food production. Additionally, more studies are needed to understand the effect of antibiotics on realistic, field scale, conditions to fully understand the impact on environmental and human health.

**Importance:** Agricultural soils are increasingly contaminated with complex mixtures of antibiotics from various biogenic sources, yet we lack a clear understanding of their specific ecological impact. While many studies investigate antibiotics, they often are studied in pollution sources like manure which contain confounding factors like heavy metals. To provide clear mechanistic insight, we investigated the effects of a complex, five-antibiotic mixture on the soil-plant system, independent of other contaminants. This revealed that the effect of antibiotics extends beyond selecting for antibiotic resistance. Specifically, the reduction of prokaryotic diversity and plant growth-promoting bacteria under antibiotic exposure can have potential detrimental effects on plant and soil health. Moreover, we found that antibiotic exposure can reduce plant biomass and nitrogen uptake, but this is highly plant dependent. This research highlights the critical need to monitor antibiotic pollution due to its potential detrimental effect on plant health and alterations to the soil microbiome.

## 1. Introduction

There is a growing need for organic fertilizers to support not only food production, but also environmental sustainability (Gamage et al., 2023). It is increasingly recognized that recycling biogenic wastes like animal manure and human excreta into organic fertilizer is a way to increase fertilizer accessibility (Trimmer et al., 2017), soil health (Urra et al., 2019) and plant growth (Luo et al., 2018). However, these recycled organic fertilizers have also been shown to contain contaminants such as antibiotics, microplastics and heavy metals (Bünemann et al., 2024; van den Broek et al., 2024). Due to the excessive use of antibiotics in human medicine and animal husbandry, antibiotics from animal manure and sewage sludge have been detected worldwide (Frey et al., 2022; Wu et al., 2023; Yang et al., 2021). Consequently, high concentrations of antibiotics end up in agricultural soils via fertilization with animal manure and sewage sludge (Wu et al., 2023) or through wastewater irrigation (Chen et al., 2014; Tian et al., 2022). Antibiotics consist of complex molecules with varying functional groups and can be divided into several classes based either on chemical structure (Kümmerer, 2009) or on the mode of action, such as inhibiting DNA replication or protein synthesis (Halawa et al., 2023). Tetracycline antibiotics (inhibition of protein synthesis) are most frequently detected in animal manure and biosolids and generally among the highest detected concentrations of all classes of antibiotics. Other frequently detected antibiotics classes in organic fertilizers are fluoroquinolones (inhibition of DNA synthesis), sulfonamides (inhibition of folic acid synthesis) and macrolides (inhibition of protein synthesis) (Cycoń et al., 2019; Frey et al., 2022; Pan and Chu, 2016). The detected antibiotic concentrations generally remain below 0.1 mg kg^−1^ range in agricultural soils worldwide, while occasionally higher concentrations have been reported, for example 0.37 mg kg^−1^ fluoroquinolone in Austria and 0.3 mg kg^−1^ sulfonamide in the UK (Frey et al., 2022)

It has been found that once antibiotics enter the soil, they can decrease microbial diversity (Cleary et al., 2016), change soil microbial community structure and lower microbial biomass (Chen et al., 2023). In addition, while antibiotics can suppress certain plant pathogens in soil such as *Pseudomonas fluorescens* and *Bacillus amyloliquefaciens* (Arseneault and Filion, 2017), they can also decrease the abundance of plant growth-promoting (PGP) microorganisms such as *Streptomyces* and *Sulfuricaulis*, which are known to stimulate crop growth and improve crop health (Ren et al., 2024). Moreover, exposure to the sulfonamide sulfadiazine has been shown to increase microbial genera harboring pathogens like *Clostridium* and *Gemmatimonas* and lowering bacterial genera associated with higher soil quality such as *Lysobacter* and *Adhaeribacter* (Ding et al., 2014). As a result, there is growing concern that these changes may negatively affect soil nutrient cycling and crop health. Moreover, antibiotics can be directly phytotoxic for crops by damaging photosynthesis processes (Krupka et al., 2022), reducing nutrient uptake (Minden et al., 2017) and causing oxidative stress (L. Li et al., 2023), thereby delaying germination (Minden et al., 2017), reducing crop biomass (Carballo et al., 2022) and lowering crop quality by reducing nitrogen (N) uptake (Fiaz et al., 2023).

In addition to impacts on soil and plant health, antibiotic exposure in agricultural systems following fertilizer application has been linked to increased presence of antibiotic-resistant bacteria (ARB) (Wu et al., 2023). While ARB occur naturally even in pristine environments due to resource competition among microorganisms (Larsson and Flach, 2022), they are much more prevalent in environments with increased anthropogenic influence (Gattinger et al., 2024). ARB contain antibiotic resistance genes (ARGs) encoding the resistance mechanism to a specific antibiotic or a group of antibiotics such as enzymatic activation proteins that break down the antibiotic or efflux pumps that transport the antibiotics outside the microbial cell. Mobile genetic elements (MGEs) can make ARGs mobile either within a genome, or across genomes. ARGs associated with MGEs can then move through a population using horizontal gene transfer (HGT) (Larsson and Flach, 2022) making MGEs an important driver of antibiotic resistance (Delgado-Baquerizo et al., 2022). Very high concentrations of antibiotics (≥10 mg kg^−1^) can be found in manure but are rarely detected in soil even when directly treated with manure (Cycoń et al., 2019; Frey et al., 2022). Consequently, the concentration of antibiotics is often lower than the minimal inhibitory concentration (Cycoń et al., 2019; Larsson and Flach, 2022). Bacteria may not be killed outright at these sub-inhibitory levels; instead, this low concentration can facilitate the development of resistance through increased mutation rates and enhanced HGT (Laureti et al., 2013). Conversely, high doses of antibiotics exert strong selective pressure, killing susceptible bacteria and inadvertently leaving an ecological niche open. This allows resistant bacteria to rapidly thrive and proliferate. This process has been extensively described in gut microbiome studies (Buffie and Pamer, 2013), and similarly, soil studies show that both low and high concentrations of antibiotics can increase ARGs (Lau et al., 2020; Ren et al., 2024). Transfer of environmental ARGs to clinical events is rare, but can have severe consequences for human health (Larsson and Flach, 2022; Waglechner and Wright, 2017). One way this transfer can happen is through consuming (raw) produce (Zhou et al., 2020), and ARB and ARGs have been found on crops like radish (Ransirini et al., 2024) and spinach (Richter et al., 2020) which are often eaten raw. Therefore, the increased antibiotic exposure of soil microorganisms through recycled fertilizers and reclaimed water application is a risk for human health.

Considering the negative environmental consequences associated with antibiotic pollution, it is vital to understand how antibiotics affect the soil microbial community as this may ultimately impact crop yields as well as environmental and human health. To date, research into the impact of antibiotics on the soil microbial community has largely focused on the direct impact of contamination sources (i.e. manure, sewage sludge). While these studies are important to understand real-life scenarios of environmental antibiotic dissemination, they often introduce not just antibiotics, but also heavy metals and nutrients to the soil, thereby confounding the observed effects on both soil microbial communities and their antibiotic resistance. To better understand antibiotic-specific microbial responses with limited influence of complex environmental variables, mechanistic investigations under controlled conditions are necessary. Moreover, mechanistic studies often investigate at most three antibiotics at the same time (Chen et al., 2023; Cleary et al., 2016; Lau et al., 2020), while agricultural soils can contain many antibiotics at the same time (Wu et al., 2023). **The goal of this study therefore was to understand the impacts of five co-applied antibiotics on the soil prokaryotic and fungal community diversity and composition, ARGs and MGEs, and plant health.** To simulate the exposure of antibiotics in organic fertilizers to the soil microbial community, we selected five structurally different antibiotics from different classes: the tetracycline chlortetracycline (CTC), the fluoroquinolone enrofloxacin (ENR), the macrolide clarithromycin (CLR), the sulfonamide sulfamethoxazole (SMX) and the diaminopyrimidine trimethoprim (TMP) which is often prescribed together with SMX and inhibits folic acid synthesis. These five antibiotics were co-applied to soil at varying concentrations of 0, 0.1, 1 and 10 mg kg^−1^ soil dry weight (DW) for each antibiotic equally. Their effects on soil microbial community structure, ARG and MGE abundances, plant growth and plant N uptake were studied in a greenhouse trial where two crop species, spinach (*Spinacia oleracea*) and radish (*Raphanus sativus*), were grown to maturity for a total of six weeks. The effects of antibiotic addition to plant growth and nutrient uptake, soil microbial community diversity and taxa, ARGs and MGEs were studied in soil samples after six weeks of exposure. **We hypothesized that the increasing antibiotic concentrations would change the soil-microbe-plant system by (i) reducing microbial α-diversity and altering microbial β-diversity, with a stronger effect on the prokaryotic community compared to the fungal community (ii) changing the microbial taxonomic composition, leading to reduction of PGP microorganisms, while increasing antibiotic-resistant microorganisms (iii) increasing ARGs and MGEs and (iv) decreasing plant productivity.**

## 2. Methods

### 2.1. Greenhouse trial and sample collection

Swiss topsoil (0-20 cm), free from the selected antibiotics and with a pH of 7.4, total organic carbon of 2.6% and total N of 0.23%, was used in the experiment. Further details on the soil properties are provided in Table S1. The soil was dry-sieved to Ø 5 mm prior use. Antibiotics used in this study were chosen to represent a range of physicochemical characteristics, for example with molecular weight from 253 (SMX) to 748 g mol^−1^ (CLR), water solubilities from 1.7 (CLR) to > 600 mg L^−1^ (SMX, CTC) and consisting of different functional groups (Avisar et al., 2010; Boxall et al., 2006; Cycoń et al., 2019; Lin and Gan, 2011; McFarland et al., 1997; Nowara et al., 1997; Sarmah et al., 2006; Stephens, 1956; Stoob et al., 2007). Further details on the chemicals used for the antibiotic analysis and physicochemical characteristics of the antibiotics are provided in Table S2 and Table S3. Antibiotics were added to the soil to reach nominal individual antibiotic concentrations of 0.1, 1, and 10 mg kg^−1^ soil DW (further referred to as c0.1, c1, c10). The stock solutions were prepared individually in methanol (TMP, CTC, ENR) or acetone (SMX, CLR) and added to autoclaved sand (500 g). A corresponding volume of pure solvents was added to the control units without antibiotic addition (namely 4.5 ml of MeOH and 2 ml acetone) (antibiotic concentration 0 mg kg^−1^, further referred to as c0). The sand was mixed thoroughly by shaking in a plastic bag. The bags were left open for 30 minutes after spiking to allow the solvents to evaporate and then pre-weighed soil (950 g) was added to the sand and mixed thoroughly. Sand amendment was used to ensure homogeneous antibiotic mixing and inhibit soil compaction during the experiment (soil:sand ratio 3:1 vol:vol). To verify that the spiking was successful and to confirm the antibiotic concentrations at D0, additional pots for each treatment were prepared (n=6). The additional pots were spiked and prepared accordingly, sampled directly (D0) by collecting 150 g of the soil-sand mixture and stored at -20 °C until analysis.

To establish the experimental replicates, the spiked soil-sand mixture was transferred to pots (Ø 15 cm), and three pre-germinated spinach (*Spinacia oleracea*) (48h, 20℃, in dark) or radish (*Raphanus sativus*) (12 h, 20℃, in dark) seeds were planted in each pot, with six replicates of each antibiotic concentration and plant treatment combination. Pots were watered to 80% water holding capacity and irrigated every second day throughout the experiment. A greenhouse trial was conducted at controlled climatic conditions (set temperature 20 ⁰C, range 19-24 ⁰C, 16:8 light: dark) and the duration of the experiment was set to 6 weeks. The experiment followed a randomized complete block design (Dean et al., 2015), where experimental units were divided into 6 blocks, each containing one replicate of each treatment. The experiment was established over the course of three consecutive days, where two blocks were started each day. The positions of the pots within the blocks and the block’s position in the greenhouse were randomized weekly. After six weeks (D42), the plants were harvested, the above- and belowground biomass were weighed separately, and the length of roots and shoots were measured. The bulk sand-soil mixture was gently homogenized prior to sample collection. Two separate bulk sand-soil mixture samples were collected; 50 g for microbial analyses and 300 g for determination of soil total antibiotic concentrations and soil pH. The plant and soil samples were stored at -20 °C immediately after collection until further processing.

### 2.2. Antibiotic analysis

The total antibiotic concentration of the bulk sand-soil mixture was determined following the method described in Shi et al. (2022), with the exception that the extract cleanup with dispersive solid-phase extraction was not conducted, as it was found not to be required after preliminary testing (results not shown). The acidity-regulated extraction-partition-concentration protocol by Shi et al. (2022) is based on sample extraction with solvents (acetonitrile, acidified with 5% formic acid and potassium phosphate buffer, pH 3). The extraction method and chemicals used in the extraction are described in further details in the supplementary information (Supplementary Text 1). The bulk sand-soil samples were extracted directly after thawing. Here, approximately 2.5g (c0, c0.1, c1) or 1.25g (c10) of moist sample was extracted, the determined total concentrations were calculated based on exact sample masses and corrected with pre-determined dry weight content of the sample (80.5 ± 1.6%) and soil content in the sand-soil mixture (65% DW). Extracted samples were analyzed on an Agilent liquid chromatography-triple quadrupole mass spectrometry system (LC-MS/MS, 6470, Agilent Technologies). The absolute recoveries ranged from 77.5 % (SMX) to 104.8 % (CLR) and the determined limits of quantifications (LOQs) were ≤ 3.19 µg kg^−1^ DW with highest LOQ observed with CTC (Table S5). Further details on analytical method, and instrument and method precision are provided in the supplementary information (Supplementary Text 2 and 3).

### 2.3. Plant nutrient and soil pH analysis

A subsample of the frozen radish roots (belowground) and leaves (aboveground), as well as spinach leaves (not enough material collected for spinach root assessment) was dried (40°C, 7 days) were ground into a fine powder, homogenized and subsampled to measure nitrogen (N) and water content. The biomass of spinach roots was too small to sample for analysis, and therefore only the spinach leaf (aboveground) samples were processed. Total moisture content for each plant was determined gravimetrically and used to calculate the dry weight biomass for each plant. The dried plant material was used to assess the total C and N content via dry combustion (LECO CHN628 Series Elemental Determinator) with isotope ratio mass spectrometry (IRMS) using the Elemental Analyzer Thermocycler Flash IRMS (EA IsoLink CN) and IRMS Delta V Plus Isotope Ratio MS (Thermo Fisher Scientific, Switzerland). Two different plant standards (124 *Medicago sativum* (44.8% C, 2.8% N) and 172 *Prunus laurocerasus* (48.0% C and 1.2% N) (International Plant-Analytical Exchange, Wageningen Evaluation Programs for Analytical Laboratories) were included in the analysis to ensure the validity of the results. In case the sample mass of individual replicate was too small for sample analysis, the missing values of C and N in roots (radish only), and leaves (radish and spinach) were filled using mean replicate values (radish roots: c1 one missing value, c10 four missing values, radish leaves: c1 one missing value, c10 three missing values, spinach leaves: c1 one missing value, c10 one missing value). Soil pH was determined in a 0.01 M CaCl_2_ solution using a pH meter (713 pH Meter, Metrohm, Switzerland).

### 2.4. DNA extraction and sequencing analysis

For metabarcoding, DNA extraction of sand-soil samples (250 ± 2 mg) was performed in randomized order with the DNeasy PowerSoil Pro Kit according to the manufacturer’s instructions using the QIACube Connect System (Qiagen, Hilden, Germany). PCR was conducted on the normalized samples targeting the 16S rRNA gene (V4 region) with 341F and 806R primers (Frey et al., 2016) and the ribosomal ITS region with ITS3ngs and ITS4ngs primers (Tedersoo and Lindahl, 2016) (Microsynth, Balgach, Switzerland) using TRUESEQ sequencing tags (Illumina, San Diego CA, USA) (See Table S6 for the primer sequences and Supplementary Text 5 for more details on the PCR protocol). The triplicate PCR products were pooled and sent to the Functional Genomics Center Zurich (FGCZ, Zurich, Switzerland) for indexing and sequencing. Indexed PCR products were purified, quantified, and pooled in equimolar ratios before pre-sequencing on the Illumina NextSeq 2000 platform (Illumina) to inform library re-pooling for optimal equimolarity across samples. Final sequencing was conducted using the v3 chemistry (PE300) on the Illumina NextSeq 2000 platform (Illumina).

### 2.5. Analyses of soil prokaryotic and fungal community

A customized bioinformatics pipeline was used to process the sequencing data as previously described (Longepierre et al., 2021). Briefly, the sequence data quality was inspected using fastqc (Andrews, 2010) and VSEARCH (Rognes et al., 2016). PhiX was removed using Bowtie2 (Langmead and Salzberg, 2012), primers were trimmed using cutadapt (Martin, 2011) and polyG tails were removed with fastp (Chen et al., 2018). Forward and reverse reads were merged (minimum merge length 300, quality truncated at phred score 7) using the fastq_mergepairs function in VSEARCH and low quality reads (expected error > 1) were filtered using the fastq_filter function in VSEARCH. Filtered reads were dereplicated, delineated and chimeras were removed using the functions derep_fulllength, cluster_unoise which uses the UNOISE algorithm (R. C. Edgar, 2016a) and uchime3_denovo which uses the UCHIME algorithm (R. C. Edgar, 2016b) in VSEARCH. Metaxa2 (16S rRNA gene) (Bengtsson-Palme et al., 2015) and ITSx (ITS2 region) (Bengtsson-Palme et al., 2013) were used to verify the target region. SILVA version 138.1 (Pruesse et al., 2007) and UNITE version 9.0 (Abarenkov et al., 2024) databases were trimmed to match the target region spanned by the primers using cutadapt, and taxonomy was assigned using the sintax function sintax algorithm (R. Edgar, 2016) in VSEARCH using a cutoff of 70%. The raw sequences were deposited in the European Nucleotide Archive with accession number PRJEB95045.

### 2.6. Database screening for pathogens and plant-beneficial microorganisms

Relevant pathogens and plant-beneficial bacteria were identified in our samples by comparing the taxonomic classifications to multiple databases. Bacterial human pathogens were identified at species level using a list of 1513 infectious bacterial pathogens composed by Bartlett et al. (2022). Bacterial plant pathogens and plant-beneficial bacteria were identified at genus level using the plant-beneficial bacteria (PBB) database containing 398 genera and the Phytopathogen database containing 258 species collected by Li et al. (2023). The database by Li et al. (2023) further specifies the PBB into broad categories such as biocontrol and stress resistance and subcategories such as N fixation and siderophore production. Fungal pathogens (animal or plant pathogen) and PGP fungi (endophytic, epiphytic and saprotrophic fungi) were identified using FUNGuild (Nguyen et al., 2016). For the FUNGuild database categorization, pathogens and parasites were classified as animal or plant pathogenic, while ectomycorrhizal, endophytic, epiphytic, arbuscular mycorrhizal, ericoid and saprotroph fungi were classified as plant-beneficial. When a genus was identified as both beneficial and pathogenic according to the PBB database, it was classified on the most likely scenario based on literature. Moreover, the presence of ESKAPE organisms (*Enterococcus faecium*, *Staphylococcus aureus*, *Klebsiella pneumoniae*, *Acinetobacter baumannii*, *Pseudomonas aeruginosa* and *Enterobacter* species) was investigated as these are six highly virulent multi-resistant bacteria that are well-known pathogens causing hospital-acquired infections (Rice, 2008). The genera significantly impacted by antibiotic treatment were screened for common antibiotic resistance in The Comprehensive Antibiotic Resistance Database (CARD) database (Alcock et al., 2020) and antibiotic-related mechanisms using Web of Science using the keywords “genus” AND “antibiotic resistance”. Then they were categorized as “pollutant degrader”, “antibiotic degrader”, “antibiotic producer”, “antibiotic resistant”, “ARG carrier” or “unclassified”. Genera associated with ARGs but not confirmed carriers were categorized as unclassified.

### 2.7. PCR and quantitative PCR of ARGs and MGEs

Antibiotic resistance genes that are commonly associated with the application of organic fertilizers and are related to the resistance to the selected antibiotics were chosen for PCR and qPCR analysis, namely *sul1* (SMX resistance) (Bischel et al., 2015; Burch et al., 2017), *dfrA12* (TMP resistance) (Xie et al., 2016), *tetQ* (CTC resistance) (Liao et al., 2018) and *qnrS1* (ENR resistance) (Zalewska et al., 2021). Clarithromycin resistance was not measured because this requires detailed investigation of the 23S gene, which is different across prokaryotic genera (Liu et al., 2017) which was outside the scope of this research. Mobile genetic elements *intI1* and *intI2* were chosen because of their associated resistance to sulfonamides and trimethoprim respectively (de los Santos et al., 2021; Hansson et al., 2002). For qualitative detection of the selected ARGs and MGEs via PCR, six primer sets targeting four ARGs and two MGEs were selected based on literature (Hu et al., 2016; Kerrn et al., 2002; Luo et al., 2010; Marti and Balcázar, 2013; Rosewarne et al., 2010) (See Table S6 for primer sequences). The validity of the selected primers was confirmed with *in silico* PCR. All selected genes were downloaded from the NCBI Nucleotide database on 04.04.2023 (National Center for Biotechnology Information, 2004) and matches to the primer were tested using cutadapt and analyzed in R using the seqinr package (Charif and Lobry, 2004). Then, PCR and qPCR were conducted (see Text section 6 for more details).

### 2.8. Statistical analyses

All statistical analyses were conducted in R version 4.4.1. (R Core Team, 2021) using RStudio Version 2024.04.2+764 (RStudio Team, 2020). Visualizations were made using ggplot2 (Wickham and Chang, 2016), patchwork (Pedersen, 2025) and RColorBrewer (Neuwirth, 2011), and refined in Adobe Illustrator (Adobe Inc., 2019). Statistical analysis of metadata (soil pH, plant biomass, plant N and antibiotic concentration (D42)), was conducted by checking for normality of data distribution using the Shapiro-Wilk test from the stats R package (R Core Team, 2021), normality of residuals using the Levene’s test from the car R package (John et al., 2020) and heteroscedasticity with the Breusch-Pagan Test from the lmtest R package (Hothorn et al., 2022). An ANOVA was conducted followed by a TukeyHSD from the stats R package when the data and residuals were normally distributed and no heteroscedasticity was found. A permutational analysis of variance (PERMANOVA) using the vegan R package (Oksanen et al., 2012) followed by a pairwise comparison using the pairwiseAdonis package (Martinez Arbizu, 2020) was conducted when the data or residuals were not normally distributed, but no heteroscedasticity was found. Multiple testing correction was done with Benjamini-Hochberg False Discovery Rate (FDR) multiple testing correction using the p.adjust function from the stats R package and significance letters based on the statistical tests were determined with the multcompView R package (Graves et al., 2024).

Quality control of the sequencing data was ensured by investigating read quality (phred scores), read length distribution and sequencing depth. Changes in α-diversity (observed richness, Pielou’s evenness and Shannon diversity) and β-diversity (Bray-Curtis dissimilarity) were calculated from 100-fold iteratively subsampled and square-root transformed ASV count tables to account for differences in sequencing depth (Schloss, 2024, 2023). PERMANOVA was used to assess the influence of antibiotic treatment, plant type and their interaction on α-diversity. For β-diversity, a principal coordinates analysis (PCoA) using the stats R package was conducted (cmdscale function). The vegan R package was used to determine significant effects of plant type and metadata (soil pH, plant biomass, plant C, plant N, antibiotic concentration (D42), ARGs and MGEs) on β-diversity. First, a full Distance-based Redundancy Analysis (partial dbRDA) model was created using plant type and all metadata as response variables with the capscale function. We did not include treatment in the model here, because it would obscure the significant influence of the individual antibiotic concentrations at D42. Then, the significance of each variable within this full model was assessed using the permutest function. The significant variables were then used to make the partial dbRDA model using the capscale function, namely plant biomass, *sul1 16S*^−1^ abundance, chlortetracycline concentration (D42), enrofloxacin concentration (D42) for prokaryotes and plant type, plant biomass, *sul1 16S*^−1^ abundance and leaf N concentration for fungi. The first two constrained axes of the partial dbRDA model were visualized using an ordination plot. When assessing soil antibiotic concentrations, values below LOQ were replaced with 0. To assess the influence of antibiotic treatment, plant type and their interaction on the abundance prokaryotic and fungal taxa, individual PERMANOVA were conducted for each phylum and genus using Euclidean distance. The models included antibiotic concentration, plant type, and their interaction and the results from these analyses were corrected for multiple testing using FDR. The results were categorized as significantly impacted by antibiotic treatment when *p ≤ 0.05*.

To further classify microbial genera significantly impacted by antibiotic treatment, the relative abundance on genus level was assessed using Generalized Linear Models (GLMs). Relative abundances were modeled using a quasibinomial family with a logit link function, a common choice for proportional data which often exhibits overdispersion. A small constant (1×10^−7^) was added to all relative abundance values to handle zero observations. A linear model was fitted for each genus-plant combination (relative abundance ∼ concentration) and the statistical significance of the linear term was evaluated using Type II Wald χ2 tests from the car R package. Based on these tests, genera were categorized into three response groups: (1) monotonic increase: assigned when the linear term for concentration was statistically significant (*p < 0.05*) and positive, (2) monotonic decrease: assigned when the linear term for concentration was statistically significant (*p < 0.05*) and negative, and (3) non-monotonic: assigned when the linear term was not significant despite the genus being identified as significantly impacted by the antibiotic treatment in the initial analysis. To determine significant interactions between the ARGs and MGEs found with qPCR (ARGs and MGEs) and the soil parameters (soil pH and soil antibiotic concentrations at D42) for both plants, Spearman correlation analysis was conducted with the rcorr function of the Hmisc R package (Harrell, 2025) and corrected for multiple testing with the stats R package. Spearman’s rho was set to 0 for non-significant correlations (*p ≤ 0.05*) and the correlations were visualized using the corrplot R package (Wei and Simko, 2024).

## 3. Results

### 3.1. Soil antibiotic concentration dynamics

The analysis of the stock solution demonstrated that the concentrations were in line with the intended spike levels, except for the CLR 0.1 mg kg^−1^ treatment spike stock. For CLR, an order of magnitude lower concentration was detected due to an error in stock solution preparation, which was noted too late for correction (Supplementary Text 4 and Table S4). The lower spike concentration of CLR were consequently also reflected in the determined soil concentrations of at D0 and D42 in the range of 0.013 –0.014 mg kg^−1^ from c0.1 treatment samples (Table 1). The D0 soil concentrations of ENR, SMX, and CTC were lower than expected based on the spiked concentrations (Table S4). For example, the determined concentrations in c10 treatment soil were 8.4, 6.8 and 5.2 mg kg^−1^ respectively. Over the course of the experiment the concentrations of CTC, SMX and TMP decreased significantly, for example for SMX the measured soils concentrations at D42 were ≤ 16% of the measured concentrations at D0 samples (Table 1). On the contrary the concentrations of ENR and CLR remained largely stable, excluding the lowest ENR treatment (c0.1), where the observed soil concentration declined from 0.06 mg kg^−1^ at D0 to 0.04 mg kg^−1^ at D42. The plant type was not found to have an effect to the measured soil concentrations.

**Table 1.** Soil antibiotic concentrations in the different antibiotic treatments at the start (D0) and end of the trial (D42). Significance between measured start (D0) and measured end (D42) concentrations determined with PERMANOVA, * (p ≤ 0.05), ** (p ≤ 0.01) and *** (p ≤ 0.005). N.D. = not detected.

| Antibiotic treatment | Measured start (D0)<br>mg kg<sup>-1</sup> | Measured end (D42)<br>Radish<br>mg kg<sup>-1</sup> | Measured end (D42)<br>Spinach<br>mg kg<sup>-1</sup> |
| --- | --- | --- | --- |
| **Chlortetracycline (CTC)** |  |  |  |
| c0 | N.D. | N.D. | N.D. |
| c0.1 | 0.089 ± 0.02 | 0.027 ± 0.01*** | 0.037 ± 0.001*** |
| c1 | 0.95 ± 0.28 | 0.27 ± 0.02*** | 0.24 ± 0.01*** |
| c10 | 5.28 ± 1.02 | 1.95 ± 1.33*** | 2.02 ± 0.08*** |
| **Clarithromycin (CLR)** |  |  |  |
| c0 | N.D. | N.D. | N.D. |
| c0.1 | 0.014 ± 0.002 | 0.014 ± 0.001 | 0.013 ± 0.001 |
| c1 | 1.09 ± 0.25 | 1.16 ± 0.05 | 1.06 ± 0.10 |
| c10 | 11.33 ± 2.82 | 11.17 ± 0.94 | 11.23 ± 0.59 |
| **Enrofloxacin (ENR)** |  |  |  |
| c0 | N.D. | N.D. | N.D. |
| c0.1 | 0.062 ± 0.006 | 0.037 ± 0.002*** | 0.037 ± 0.002*** |
| c1 | 0.58 ± 0.15 | 0.53 ± 0.05 | 0.53 ± 0.03 |
| c10 | 8.43 ± 1.57 | 8.86 ± 0.86 | 9.01 ± 0.66 |
| **Sulfamethoxazole (SMX)** |  |  |  |
| c0 | N.D. | N.D. | N.D. |
| c0.1 | 0.041 ± 0.005 | 0.007 ± 0.001*** | 0.005 ± 0.001*** |
| c1 | 0.49 ± 0.11 | 0.074 ± 0.02*** | 0.067 ± 0.02*** |
| c10 | 6.79 ± 1.36 | 1.07 ± 0.21*** | 1.092 ± 0.09*** |
| **Trimethoprim (TMP)** |  |  |  |
| c0 | N.D. | N.D. | N.D. |
| c0.1 | $0.13 \pm 0.02$ | $0.004 \pm 0.0004^{***}$ | $0.004 \pm 0.001^{***}$ |
| c1 | $1.12 \pm 0.22$ | $0.043 \pm 0.004^{***}$ | $0.041 \pm 0.006^{***}$ |
| c10 | $10.79 \pm 2.20$ | $4.82 \pm 0.47^{***}$ | $4.82 \pm 0.37^{***}$ |

### 3.2. Impact of antibiotics on the soil microbial community

In total, we found 37089 prokaryotic ASVs and 3605 fungal ASVs. Prokaryotic α-diversity was significantly influenced by antibiotic treatments and plant type as determined by pairwise PERMANOVA. Observed richness, Pielou’s evenness, Shannon diversity index and Inverse Simpson diversity index decreased with increasing antibiotic concentration (Fig. 1, Table S8). For the fungal community, there were no significant effects for both antibiotic treatment and plant type, except for evenness which significantly increased for radish at c1 compared to c0 (Fig. 1D). Observed richness, Shannon diversity and Inverse Simpson were much higher for prokaryotic communities than for fungal communities (Table S8). Prokaryotic β-diversity was significantly affected by antibiotic treatment (*p = 0.001*) but not plant type (*p = 0.457*). β-diversity did not significantly differ between the c0 and c0.1 treatments, whereas all other pairwise comparisons between treatments showed significant differences (*p = 0.006*) (Fig. S1A, Table S9). Further investigation using partial dbRDA indicated that antibiotics CTC (*p = 0.0038*) and ENR (*p = 0.005*), as well as *sul1* gene abundance (*p = 0.0007*) and plant biomass (*p = 0.0001*) significantly influenced prokaryotic β-diversity (Fig. 1C). Fungal β-diversity was significantly affected by both antibiotic treatment (*p = 0.001*) and plant type (*p = 0.004*) for the fungal community. Within the antibiotic treatments, only c10 was significantly different from the other antibiotic treatments (*p = 0.006*) (Fig. S1B, Table S9). Partial dbRDA for fungal β-diversity revealed significant correlations with plant type (*p = 0.0009*), total plant biomass (*p = 0.0001*), N leaf concentration (*p = 0.0078*) and *sul1* gene abundance (*p = 0.0006*) (Fig. 1F).

**Figure 1.**
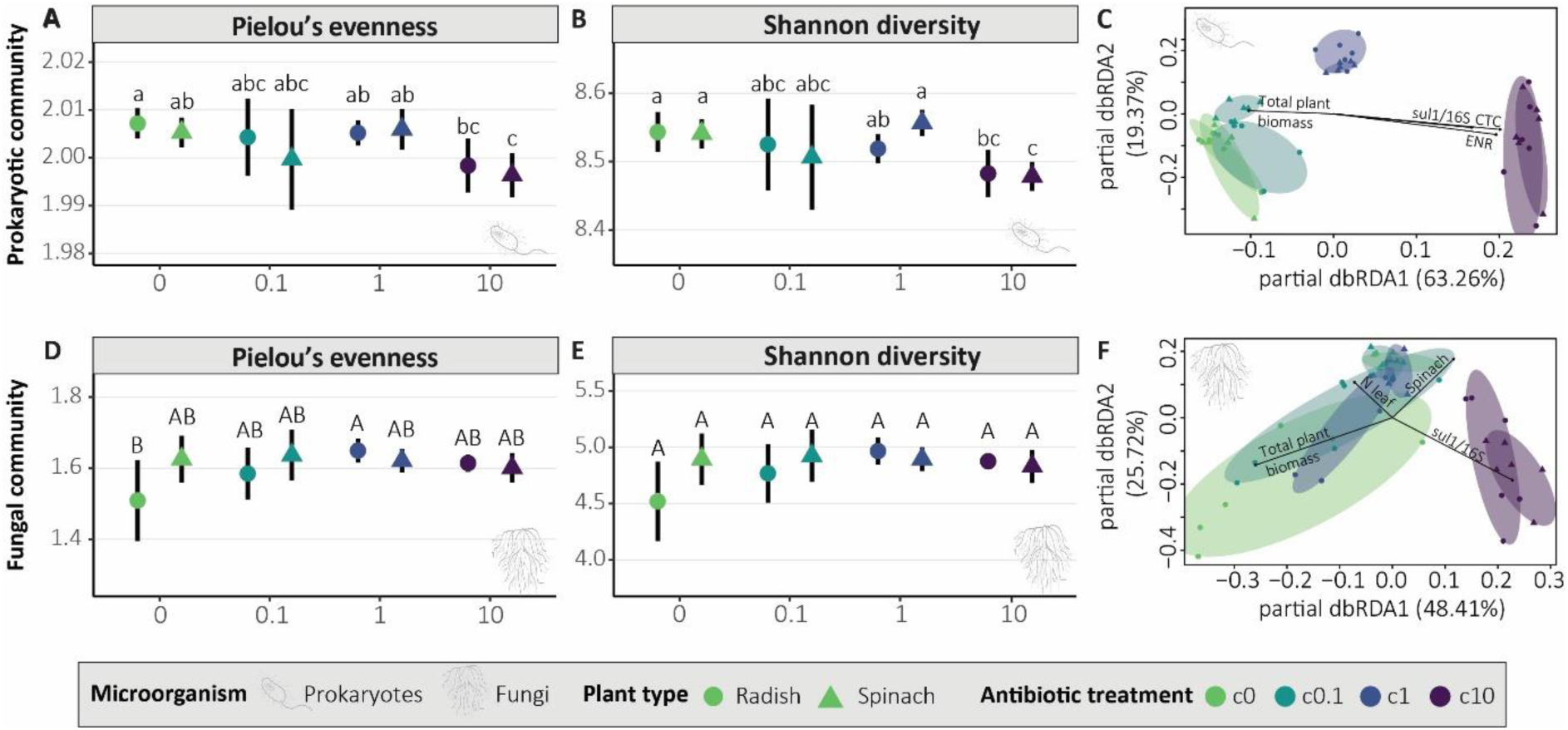
Alpha- and beta-diversity of soil prokaryotic and fungal communities. Differences in lowercase letters and uppercase letters indicate significant differences between plant type and antibiotic treatments for prokaryotic and fungal communities respectively (p ≤ 0.05, see also Table S8) (A, B, D, E). Arrows show which plant and soil properties significantly impacted β-diversity (p ≤ 0.05) (C, F). CTC = chlortetracycline and ENR = enrofloxacin (C, F).

For the prokaryotic community, 13 out of 47 phyla (27.7 %) were significantly impacted by antibiotic treatment (*p ≤ 0.05*), four of which are in the 12 most abundant phyla namely Chloroflexi (now named Chloroflexota) (*p = 0.005*), Planctomycetota (*p = 0.005*), Bacteroidota (*p = 0.005*) and Gemmatimonadota (*p = 0.005*) (Fig. 2A). At the genus level, 139 out of 671 genera (20.7 %) were significantly impacted by antibiotic treatment, affecting 6 of 12 most abundant genera, namely *Pir4 lineage*, *Nocardioides*, *Pirellula*, *Spingomonas*, *Solirubrobacter* and *Hyphomicrobium* (Fig. 2B). For the fungal community, 6 out of 15 phyla (37.5 %) were significantly affected by antibiotic treatment, namely Ascomycota, Aphelidiomycota, Mortierellomycota, Basidiobolomycota, Blastocladiomycota and Olpidiomycota (Fig. 2C) and 0 out of 447 genera (Fig. 2D).

**Figure 2.**
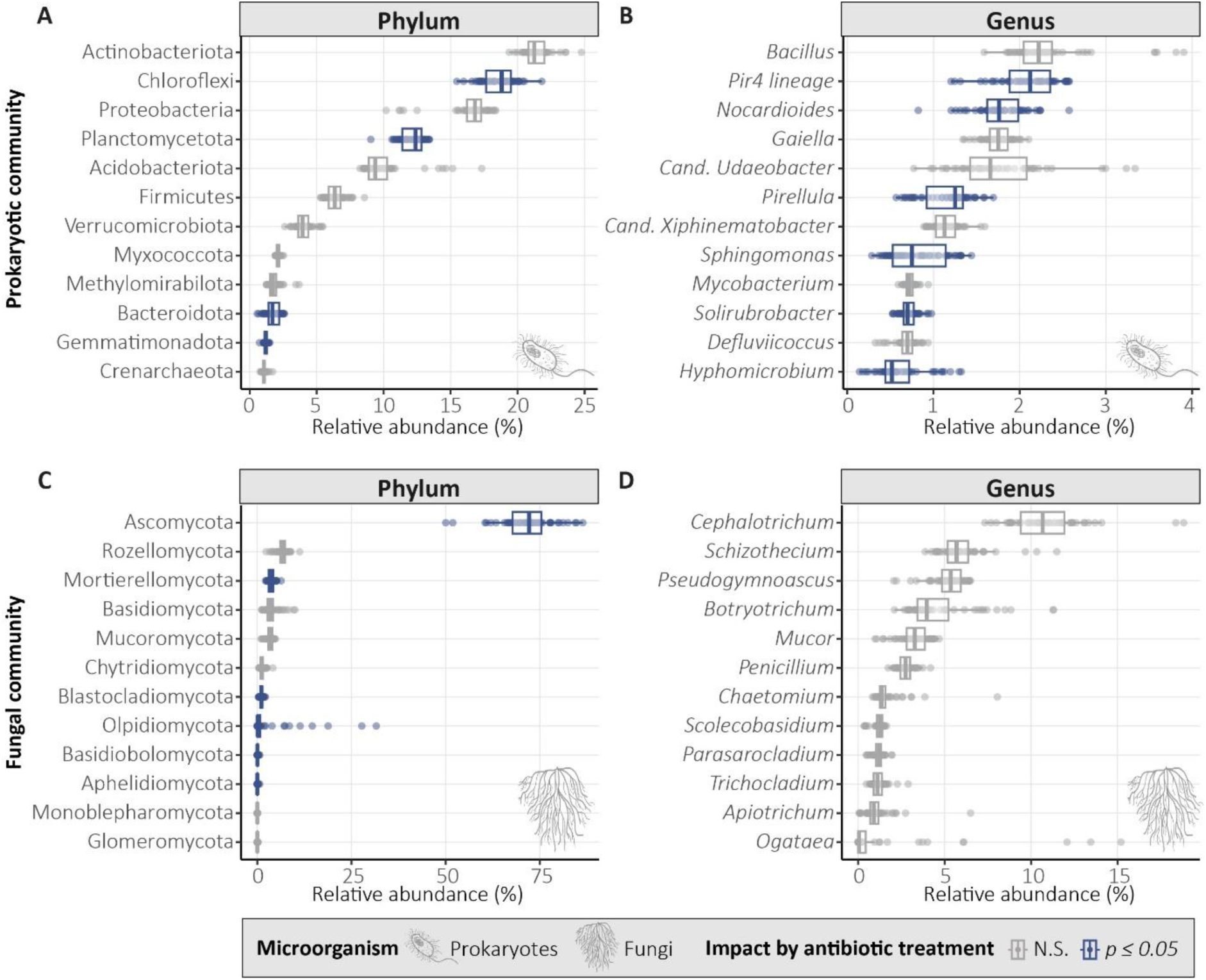
Taxonomic distribution (phyla and genera) of the 12 most abundant prokaryotic phyla (A) and genera (B) and fungal phyla (C) and genera (D). Taxonomic groups significantly impacted by antibiotic treatment (PERMANOVA, p ≤ 0.05) are colored in blue.

### 3.3. Impact of antibiotics on microbial genera and their characteristics

For prokaryotes, 65 (radish, 46.8 %) and 64 (spinach, 46.0 %) of the 139 genera followed a monotonic decrease, 20 (radish 14.4 %) and 30 (spinach, 21.6 %) genera followed a monotonic increase, 54 (radish, 38.8 %) and 45 (spinach, 32.4 %) genera followed a non-monotonic pattern (Fig. 3). Some of the monotonically decreasing genera included known phosphorus-solubilizing genera like *Adhaeribacter* and *Flavisolibacter*, indole-3-acetic acid (IAA) producing genera like *Aeromicrobium*, *Agromyces and Cupriavidus,* N-fixing genera like *Noviherbaspirillum, Pontibacter* and *Spinghomonas* and potentially human pathogenic *Brevundimonas* (Fig. 4A). Monotonically increasing genera included known IAA-producing *Spinghobium*, siderophores producing *Skermanella* and PGP *Jiangella*, *Polaromonas* and

**Figure 3.**
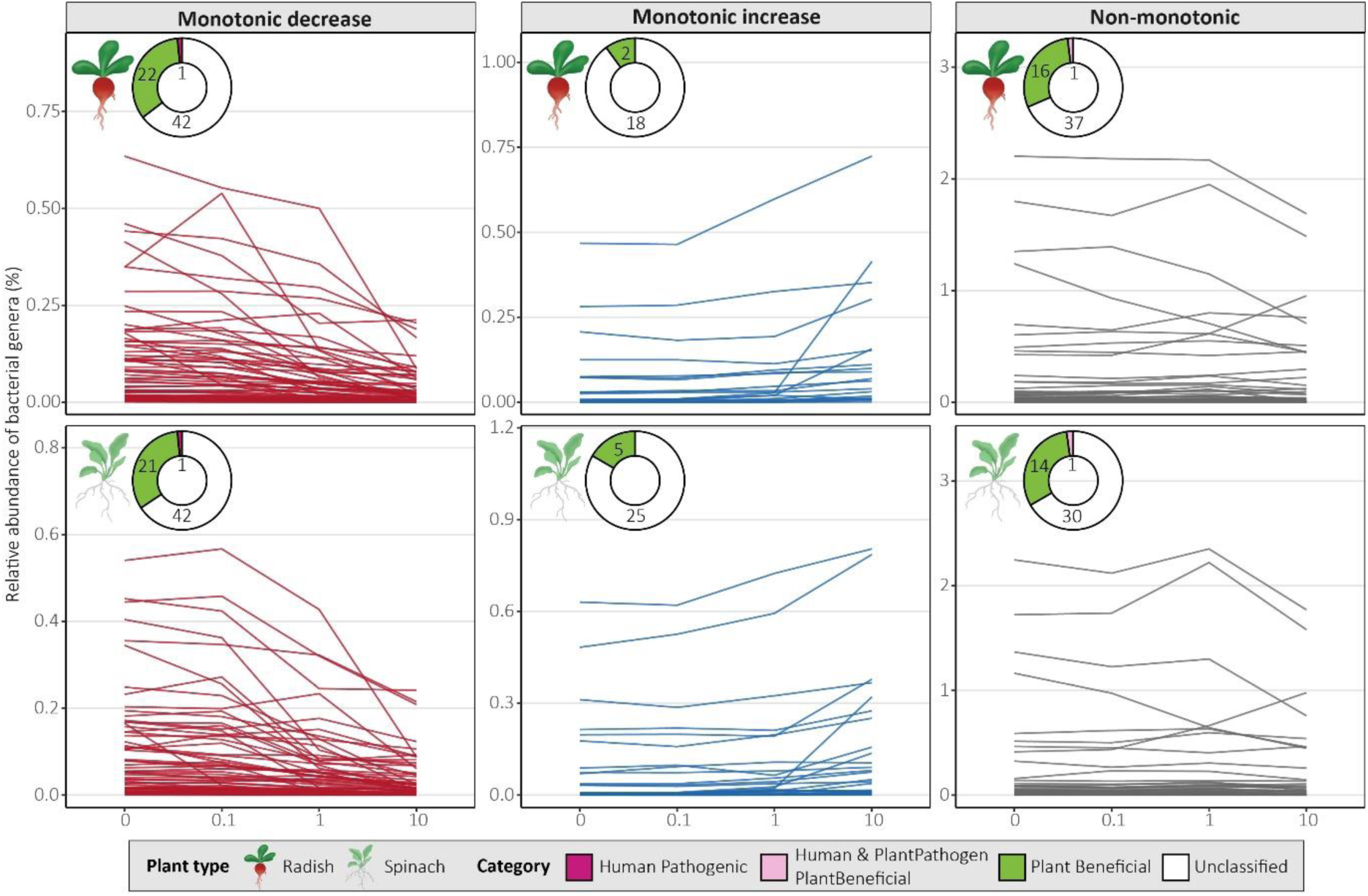
Prokaryotic genera significantly influenced by antibiotic treatment categorized by monotonic response, with each line representing a genus. Potential microbial categories are classified at genus level based on the human pathogenic bacteria database (Bartlett et al., 2022), the plant-beneficial and pathogenic bacteria database (P. Li et al., 2023) – prokaryotes that were not in these databases were labelled as ‘unclassified’. See also supplementary materials Fig. S2 for the responses of specific genera and their characteristics.

**Figure 4.**
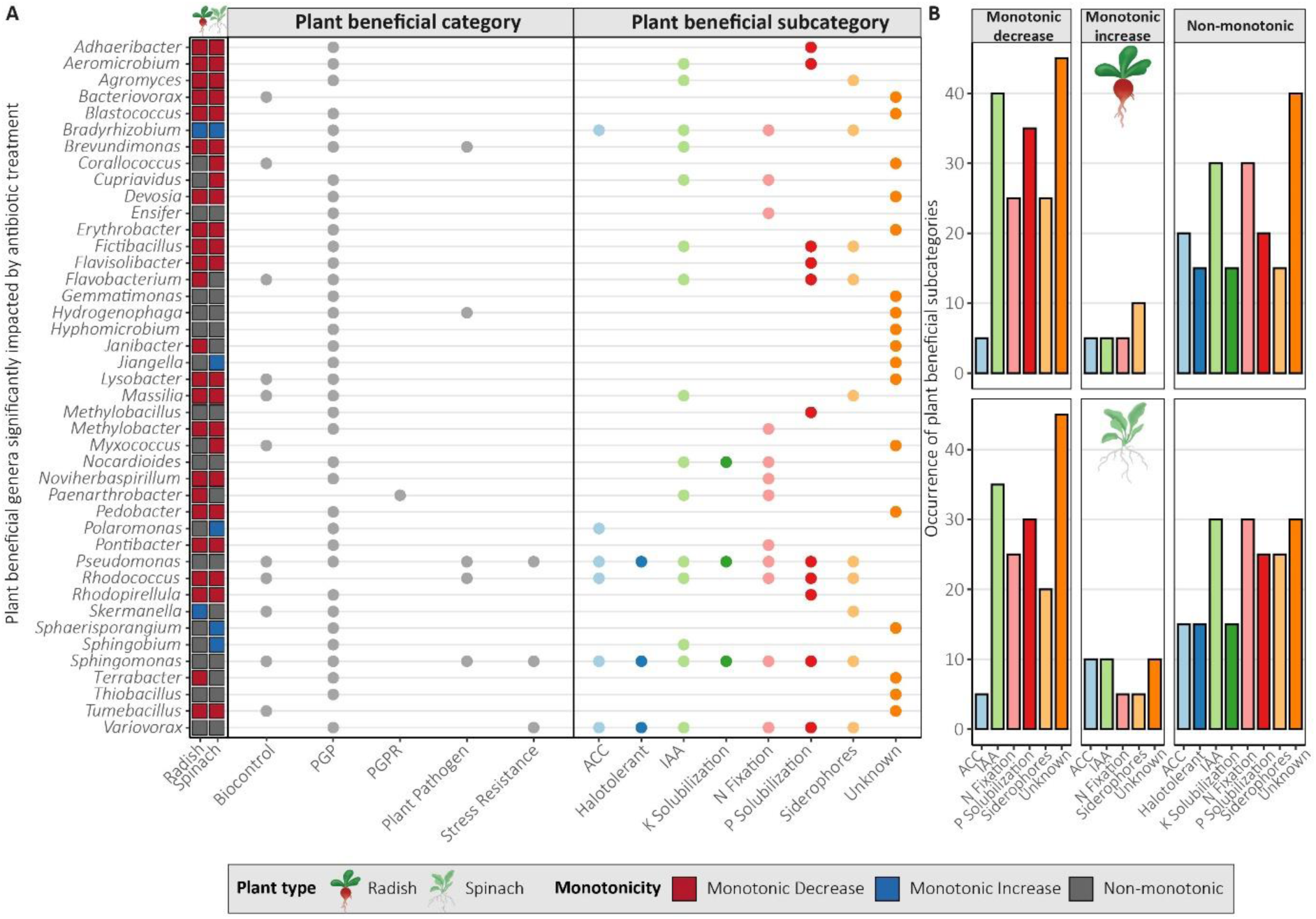
Plant beneficial categories of plant growth promoting genera significantly impacted by antibiotic treatment (A) and the distribution of the subcategories of these genera by monotonicity group (linear monotonic decrease, linear monotonic increase and non-monotonic) and plant type (B). Plant beneficial categories are based on the plant-beneficial and pathogenic bacteria database (P. Li et al., 2023). Genera that were in the database but did not have a subcategory were categorized as unknown. PGP = plant growth-promoting, PGPR = plant growth-promoting rhizobia, ACC = aminocyclopropane-1-carboxylate deaminase and IAA = indole-3-acetic acid.

*Sphaerisporangium*. Many prokaryotic genera followed a non-monotonic pattern of which many were plant-beneficial such as N-fixing genera like *Ensifer*, *Pseudomonas* and *Sphingomonas*, phosphorus-solubilizing genera *Methylobacillus* and indole-3-acetic acid producing genera *Nocardioides* (Fig. 4A). Some genera followed different patterns depending on plant type but never exhibited both a monotonic decrease and increase (Fig. 4A). In general, the monotonically decreasing genera included mostly IAA-producing, phosphorus-solubilizing and N-fixing genera, while monotonically increasing had more aminocyclopropane-1-carboxylate deaminase-producing and siderophore-producing genera (Fig. 4B). As no fungal genera were significantly impacted by antibiotic treatment, they were not analyzed for monotonicity.

None of the six ESKAPE organisms (*Enterococcus faecium*, *Staphylococcus aureus*, *Klebsiella pneumoniae*, *Acinetobacter baumannii*, *Pseudomonas aeruginosa* and *Enterobacter* species) were found in any of the samples. The 139 genera significantly impacted by antibiotic treatment contained one antibiotic-producing strain, five pollutant degraders, nine antibiotic degraders, 24 antibiotic resistant strains and 11 ARG carriers (Fig. 5). Many organisms known for their antibiotic resistance, such as *Ensifer*, *Janibacter*, *Pseudomonas* and *Variovorax* (Fig. 5), did not follow a non-monotonic pattern (Fig. 5A) but increased in c1 compared to c0 (Fig. S2). Other genera following a non-monotonic pattern increased in c0.1 compared to c0, like *Aridibacter*, *Cupriavidus* and *Myxococcus* (Fig. S2).

**Figure 5.**
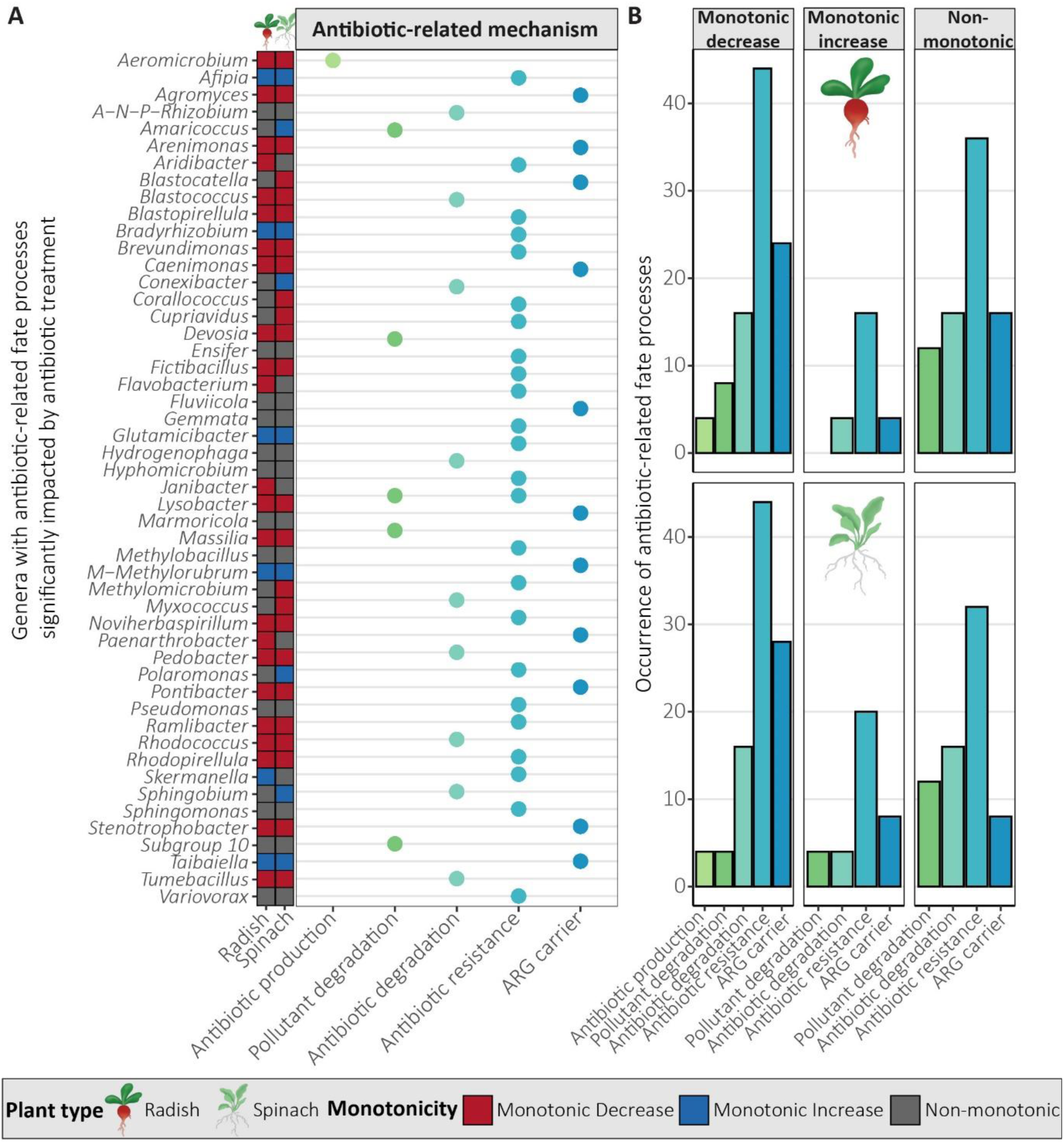
Antibiotic-related fate processes of genera significantly impacted by antibiotic treatment as determined by a literature search (Web of Science) (A) and the distribution of the antibiotic-related mechanisms of these genera by monotonicity group (linear monotonic decrease, linear monotonic increase and non-monotonic) and plant type (B). A– N–P–Rhizobium = Allorhizobium–Neorhizobium–Pararhizobium–Rhizobium and M-Methylorubrum = Methylobacter-Methylorubrum.

### 3.4. Antibiotic resistance genes and mobile genetic elements

The selected primers can successfully detect ARG *dfrA12*, *tetQ, qnrS1* and *sul1* and MGE *intI1* and *intI2* according to the *in silico* PCR analysis (Fig. S3). Both *sul1* and *intI1* were present in all samples, while *dfrA12*, *tetQ, qnrS1* and *intI2* did not amplify with the selected primers and PCR settings. *Sul1* and *intI1* abundance significantly increased in c10 compared to the other antibiotic concentrations c0, c0.1 and c1, and *sul1* also was significantly higher in c1 for spinach compared to c0 for both plants and c0.1 for spinach (Fig. S4). A correlation analysis of the genes detected, soil pH and antibiotic concentration showed that both *sul1* and *intI1* positively correlated with each other in both radish (*Spearman’s ρ = 0.85*) and spinach (*ρ = 0.54*) soils (Fig. 6). *Sul1* was positively correlated with soil pH in the spinach soil samples (*ρ = 0.58*), but not in the radish soil samples. *Sul1* and *intI1* both significantly and positively correlated with the end antibiotic concentrations, for *sul1* this was particularly strongly correlated for spinach (*ρ = 0.90 to 0.93*), and to a lesser extent in radish (*ρ = 0.65 to 0.72*). Soil pH was correlated with the antibiotic treatment for spinach and radish, where c0 showed a significantly higher pH compared to c10 for radish, while the opposite was observed for spinach (Fig. S5). The soil pH of radish and spinach became more similar with increasing antibiotic treatment concentration (Fig. S5) and soil pH was specifically lower for c0 for both radish and spinach with a soil pH of 7.02±0.06 and 6.71±0.09 respectively (Table S7), compared to starting soil pH of 7.36 before the sand was added (Table S1). Soil pH also correlated differently with the antibiotic concentrations, ranging from negative for radish (*ρ = -0.54 to -0.50*) to positive for spinach (*ρ = 0.51 to 0.59*). For the antibiotics, CTC, CTM, ENR, SMX and TMP all strongly positively correlated with each other (*ρ > 0.92*).

**Figure 6.**
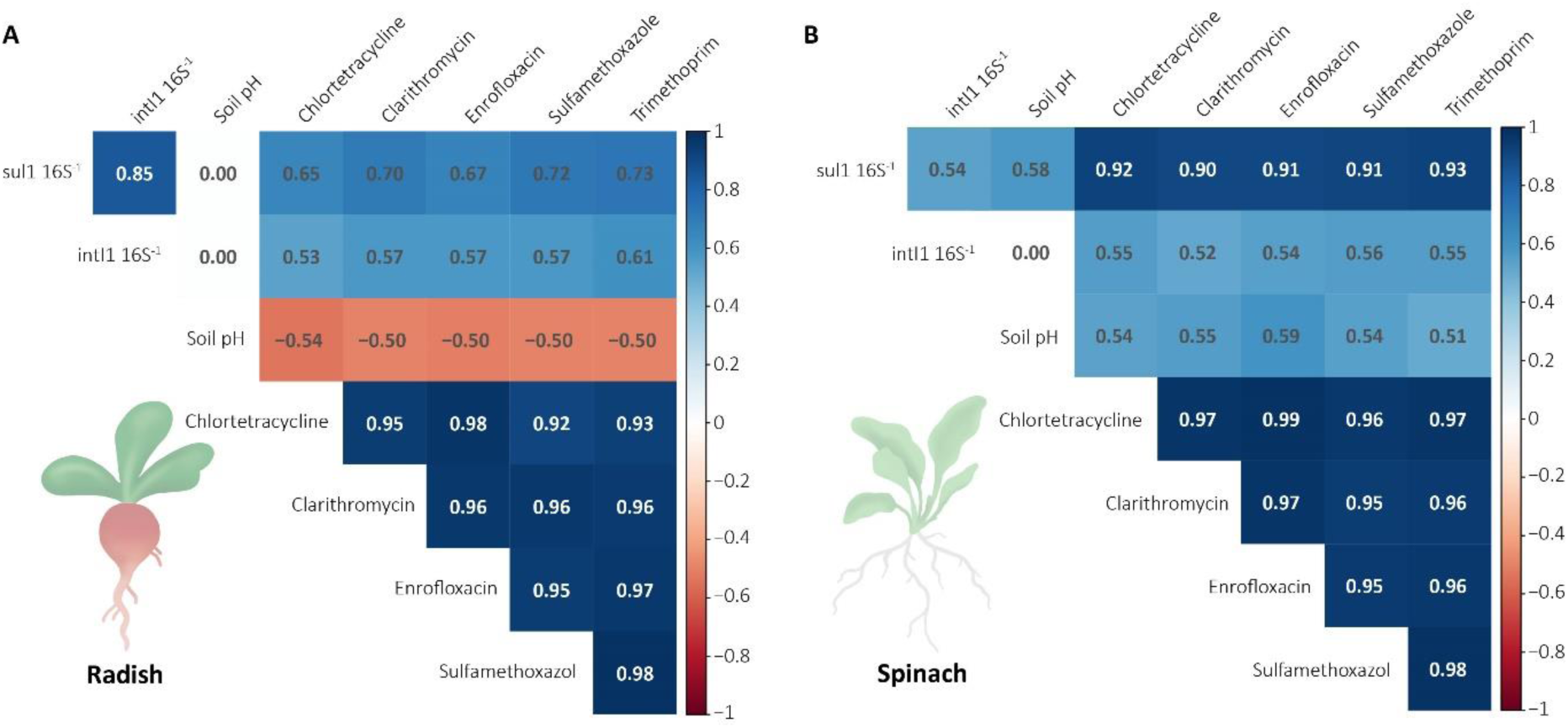
Correlation of sul1, intI1, soil pH (D42) and measured antibiotic concentrations (D42) across all treatments (c0, c0.1, c1 and c10), separated by plant type. Correlation coefficient based on Spearman’s ρ. Only significant (p ≤ 0.05) correlations are shown.

### 3.5. Impact of antibiotics on plant biomass and plant nitrogen uptake

The above- and belowground biomass of radish decreased significantly with increasing antibiotic concentrations (*p ≤ 0.05*), while spinach was not affected (Fig. 7). C10 significantly reduced aboveground N concentration for radish, while for spinach only c1 was significantly lower than the control treatment (c0). Similarly, total N uptake per plant decreased with increasing antibiotic concentrations for radish, but not for spinach (Fig. 7).

**Figure 7.**
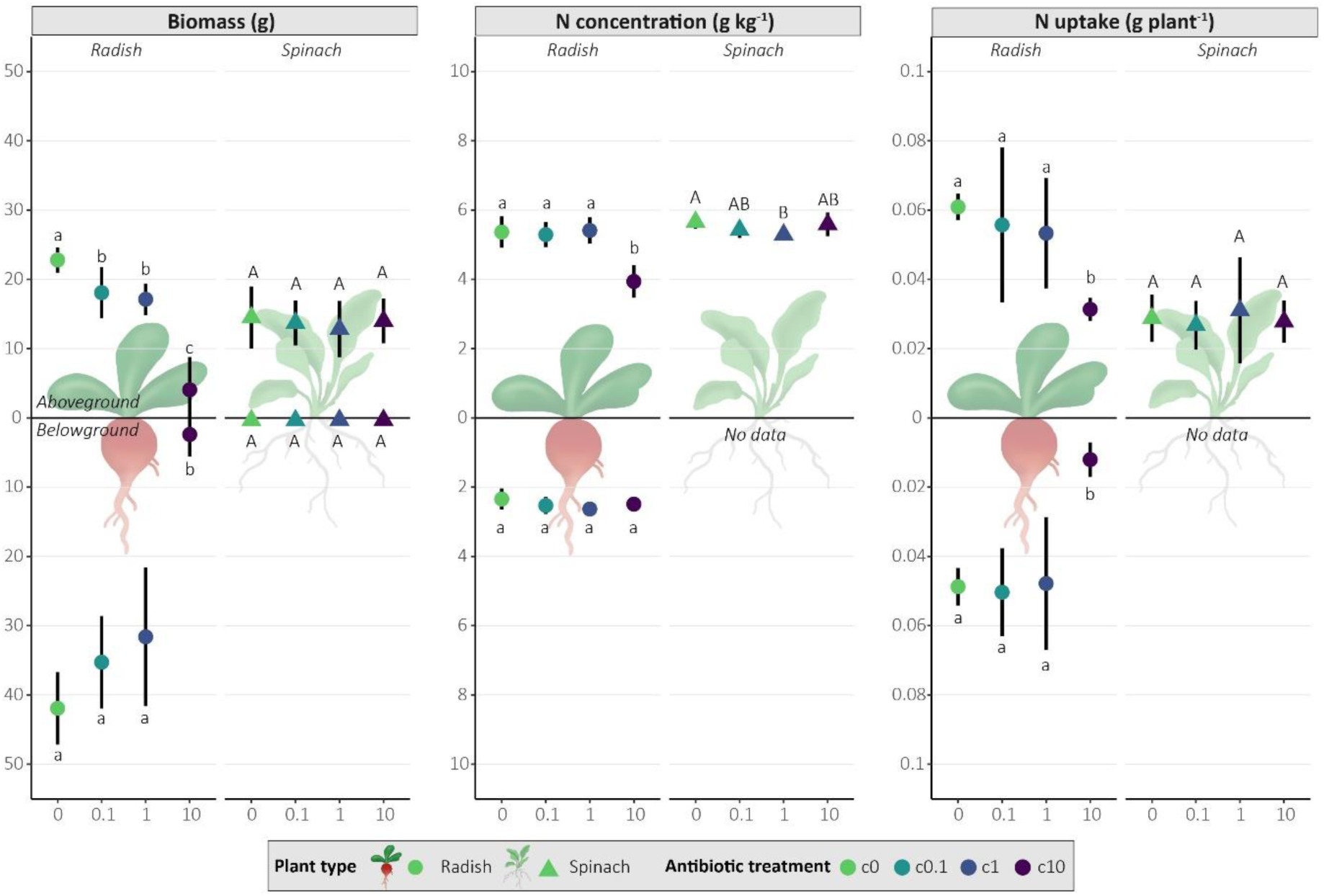
The effect of antibiotic concentration on fresh plant biomass, nitrogen (N) concentration of above- and below-ground plant tissue and total N uptake per plant in the leaves (aboveground) and roots (belowground). Differences in lower case letters indicate significant differences between radish antibiotic treatments while differences in upper case letters indicate significant differences between spinach antibiotic treatments (p ≤ 0.05, see also Table S7). Spinach root N was not measured due to insufficient biomass.

## 4. Discussion

### 4.1. Antibiotic type and concentration drive persistence in soil

The transformation and degradation of antibiotics in soil are driven largely by their molecular structure and physicochemical properties (Cycoń et al., 2019). While dissipation of antibiotics varies notably within individual studies, fluoroquinolones, macrolides, and tetracyclines were found to be characterized by long degradation half-lives in soil in a comprehensive review by Cycoń et al. (2019). In agreement with this outcome, our results showed that fluoroquinolone ENR and macrolide CLR concentrations remained stable over the course of the experiment (Table 1). However, the CTC soil concentration at D42 was 25-42% of the initial measured at D0, indicating favorable conditions for dissipation in our study (Table 1). Degradation half-lives from < 1 day to 34 days in soil has been reported in literature for CTC (Cycoń et al., 2019). The dissipation of SMX and TMP were most pronounced in our experiment, where at D42 only 12%, 14% and 16% (SMX) and 3%, 4% and 45% (TMP) of the initial measured (D0) concentration remained in c0.1, c1 and c10 treatments, respectively (Table 1). Although abiotic degradation processes and other mechanisms of losses (e.g. adsorption to pot surfaces, strong sorption to soil particles leading to the formation of non-extractable residues) cannot be ruled out (see further details in Supplementary Text 4), it can be assumed that microbial degradation is the primary contributor to the observed dissipation of the antibiotics as indicated by several studies demonstrating that microbes prominently contribute to the degradation of antibiotics in soils (Pan and Chu, 2016; Yang et al., 2009). However, the highest antibiotic concentration treatment may hinder antibiotic degradation due to the stronger inhibition of soil microbial activity (Cycoń et al., 2019; Demoling et al., 2009; Fang et al., 2016). This is reflected in the dissipation of TMP and ENR, where the dissipation for TMP was higher in c0.1 and c1 (≤ 97%) compared to c10 (≤ 52%) and the dissipation for ENR c0.1 was 40%, whereas the concentrations remained stable in c1 and c10. Co-application of antibiotics, as done in this experiment, may also lead to inhibition of soil microbial activity and microbial degradation, consequently leading to longer persistence of the antibiotics in soil (Kodešová et al., 2020) and continued selection pressure on the soil microbial community.

### 4.2. Antibiotics exert a stronger influence on prokaryotic diversity than on fungal diversity

Prokaryotic α-diversity significantly reduced with increasing antibiotic concentration (Fig. 1A-B, Table S8), in line with previous research (Gao et al., 2021). Prokaryotic β-diversity was significantly different for c1 and c10 compared to c0 and c0.1 (Table S9). Previous research showed that co-applied antibiotics impact prokaryotic β-diversity (Shen et al., 2021), but not single antibiotic application (Brown et al., 2022; Cleary et al., 2016). Prokaryotic β-diversity was mostly correlated with soil CTC and ENR concentrations, which remained in the soil at high concentrations throughout the experiment (Table 1). As expected, α- and β-diversity were more impacted by antibiotic treatment for the prokaryotic community than the fungal communities (Fig. 1, Table S8), because the five applied antibiotics target bacterial pathways and should have little direct effect on fungi. However, the direct impact of antibiotic exposure on prokaryotic communities, and the subsequent changes of abiotic and biotic properties, can impact fungal communities. This indirect effect could explain why fungal β-diversity was significantly different in c10 compared to the other antibiotic treatments (Table S9). Thus, we can accept our hypothesis that antibiotic exposure reduces prokaryotic α-diversity and fungal α-diversity, with fungal diversity impacted to a lesser extent than prokaryotic diversity. Similar, β-diversity was also impacted by antibiotic exposure, with prokaryotes again being more affected than fungi. These changes have the potential to negatively impact soil and plant health, because soil microbial diversity has been shown to be essential in providing ecosystem functions and services for human welfare as a driver of soil fertility and crop productivity (Delgado-Baquerizo et al., 2016; Romero et al., 2023) and protection against pathogen establishment (Van Elsas et al., 2012).

### 4.3. Antibiotic-induced shifts in key bacterial genera threaten microbial-mediated plant support mechanisms

Several bacterial genera were significantly impacted by antibiotic exposure, while no fungal genera were impacted. Only few bacterial genera were potentially pathogenic, namely *Brevundimonas* (monotonic decrease) and *Pseudomonas* (non-monotonic, but increased at c1), containing both pathogenic and PGP bacteria (Bull et al., 2010; Qin et al., 2022; Sah et al., 2021) (Fig. 3, Fig. S2). Most responsive PGP bacteria decreased or followed a non-monotonic pattern, potentially affecting N fixation, phosphorus solubilization, production of phytohormone IAA and plant stress-alleviating enzyme ACC capabilities (Fig. 3, Fig. 4). This suggests that antibiotic exposure may impact plant growth and development through the inhibition of microbial-mediated nutrient cycling, PGP phytohormone production and stress alleviating enzymes, which could partially explain reduced plant growth and N uptake for radish in c10 (Fig. 7). Monotonically increasing genera unique to spinach included PGP bacteria and potentially pollutant degraders (Fig. 4, Fig. 5) such as *Amaricoccus* (micropollutant degrader (Brunhoferova et al., 2022)), *Conexibacter* (lincomycin degradation (Lei et al., 2023)), *Jiangella* (PGP), *Polaromonas* (PGP), *Sphaerisporangium* (PGP) and *Sphingobium* (stress alleviation, biodegradation and bioremediation (Boss et al., 2022)) (Fig. 4, Fig. 5). These genera potentially could have contributed to protecting spinach against the effects of antibiotic exposure, thus explaining why it was less affected than radish (Fig. 7).

In our study, most PGP bacteria decreased (Fig. 3, Fig. 4) and only a few PGP bacteria increased and were known antibiotic-resistant or ARG carrier (Fig. 4, Fig. 5), despite the common antibiotic-resistant and stress-adapted nature of PGP bacteria (Hou and Kolodkin-Gal, 2020; Mahdi et al., 2022; Wash et al., 2022). This is potentially due to the selective pressure of co-applied antibiotics of different classes. Some responsive genera included tetracycline and sulfonamide degraders *Hydrophenophaga* (increased at c1 for radish) (Li et al., 2024; Tian et al., 2024), *Ramlibacter* (unaffected at c0.1 compared to c0) (Ajibade et al., 2023; T. Li et al., 2023; Zhang et al., 2022) and *Skermanella* (increased at c10) (Garbini et al., 2022) (Fig. S2) which likely enhanced CTC and SMX degradation (Table 1). However, co-application of antibiotics may inhibit microbial degradation, especially at high concentrations (section 4.1). Many significantly impacted genera contain antibiotic-resistant or ARG-carrying species, but few genera increased. Further research is needed to find out which species are antibiotic-resistant and if their ARGs convey resistance to the applied antibiotics. Moreover, it is important to consider that taxonomic shifts do not necessarily translate into altered functional capacities of the soil microbiome (Jiao et al., 2019). Based on our findings we accept our hypothesis that antibiotic exposure changes taxonomic composition, specifically by reducing PGP bacteria while not affecting fungi. We cannot definitively conclude that antibiotic-resistant microorganisms increased, as only few genera increased, and thus further analysis is necessary to confirm which species are antibiotic-resistant or ARG-carrying. The reduction of PGP bacterial genera may have significant impacts on nutrient cycling and plant protection mechanisms. Moreover, antibiotic exposure potentially impacted microbial antibiotic degradation and antibiotic resistance, though most genera responded non-monotonically. The genera increasing at low antibiotic concentrations such as c0.1 and c1 such as *Pseudomonas* are potentially the most concerning as these concentrations are more commonly detected in soils amended with organic fertilizers (Cycoń et al., 2019; Thiele-Bruhn, 2003). It is therefore especially critical to better understand impacts of antibiotics at these more common concentrations not only on microbial community structures, but also functional implications for plant productivity and soil health.

### 4.4. Dose-dependent selection for sul1 and intI1 under combined antibiotic pressure

Our study showed a significant increase in *sul1* and *intI1* abundance in the c10 treatment (Fig. S4), likely due to persistent selection pressure. SMX concentration in the c10 treatment remained sufficiently high throughout the experiment to continue selection pressure (Table 1), and both genes were strongly correlated with antibiotic concentrations at D42 (Fig. 6). *Sul1* and *intI1* were not linked to one specific type of antibiotic, as all antibiotics were significantly correlated with these genes (Fig. 6), making it difficult to determine whether one antibiotic exerts more selective pressure than another. We also found a strong correlation between *sul1* and *intI1* (Fig. 6), which is expected because *sul1* lacks a promotor and is always found with *intI1* (de los Santos et al., 2021). Additionally, *sul1* and *intI1* were significantly positively correlated with soil pH in spinach but negatively correlated in radish (Fig. 6), this is consistent with previous research which shows that soil pH (directly or indirectly) impacts ARGs and MGEs (Delgado-Baquerizo et al., 2022). In our study, the two plant species created significantly different soil pH (Fig. S5). As antibiotic dissipation did not differ between plants (Table 1), there is likely another driver of soil pH such as plant exudates which are known to change soil pH (Vives-Peris et al., 2020). A key consideration in interpreting our results is the lack of a positive control which is important in determining the success of amplification (Keenum et al., 2022). Consequently, we cannot definitively conclude that the lack of amplification for *dfrA12*, *tetQ, qnrS1* and *intI2* is a true absence or a technical limitation due to primer choice or PCR conditions. Unfortunately, the lack of a positive control is common in studies of ARGs and MGEs due to safety risks and requirements of working with cell cultures harboring ARGs. Nonetheless, we can accept our hypothesis that increasing antibiotic concentrations increase *sul1* and *intI1* abundance. Only c1 (spinach) and c10 (spinach and radish) increased *sul1* and *intI1* abundance (Fig. S4), indicating that lower concentrations may not pose a risk for increasing these genes under combined antibiotic exposure. However, sulfonamides like SMX are regularly used in hospitals as a first-line antibiotic and *sul1*-mediated resistance to SMX poses a significant clinical risk (Osose et al., 2025). Therefore, monitoring environmental ARGs is crucial to avoid transfer into clinical settings (Larsson and Flach, 2022), especially in agricultural settings where consumer crops may be a transfer vehicle for ARGs and pollution is common.

### 4.5. Radish exhibits greater susceptibility to antibiotic exposure than spinach

In our study, radish biomass and N uptake were much more affected by antibiotic treatment than spinach (Fig. 7). While some studies have reported no significant effect or even increased radish biomass under single or repeated antibiotic exposure (Cui et al., 2022; Jia et al., 2024; Yin et al., 2023), our findings align with concentration-dependent responses for radish biomass (Chung et al., 2017; Sidhu et al., 2019). Previous studies showed that oxytetracycline increased leaf catalase activity, indicative of oxidative stress (Cui et al., 2022) and can negatively affect radish subcellular structures (chloroplasts and vacuole) dependent on antibiotic type (Jia et al., 2024). The most significant above- and belowground biomass reduction for radish was observed in c10. However, c0.1 and c1, which more closely represent actual environmental concentrations, also had significantly lower aboveground biomass compared to c0. Although simultaneous presence of five antibiotics at identical concentrations is unlikely due to different degradation rates (section 4.1), the effects observed at these more realistic exposure levels are important to consider. Antibiotics up to 10 mg kg^−1^ may not affect spinach biomass (Mohy-u-Din et al., 2023; Nafees et al., 2025), but can increase oxidative stress (Nafees et al., 2025). In general, plant health response is highly dependent on applied antibiotic (Khan et al., 2022; Liu et al., 2009a) and plant type (Liu et al., 2009b; Minden et al., 2017) and combined application of antibiotics from different classes may increase toxicity (Timmerer et al., 2020). The difference in plant response may be further explained by the plant-dependent reduction of PGP bacteria following antibiotic exposure (Fig. 3, Fig. 4). This can further exacerbate harmful effects on the plants by reducing microbial protection mechanisms (such as oxidative stress protection) and microbial-mediated nutrient cycling. With this we can accept our hypothesis that increasing antibiotic concentrations decrease plant yield and N uptake for radish but reject our hypothesis for spinach. This highlights the complexity of soil-microbe-plant interactions and emphasizes the need for studies investigating the interplay between microbial activity (i.e. transcriptomics, enzymatic analyses) and plant health under antibiotic exposure.

## Acknowledgements

The authors would like to thank Valentine Okwonko and Dr. Carlton Poindexter for their help with selecting antibiotic resistance gene primers for this research, Annina Maier for their help in the greenhouse trial, Evangeline Yang for her help in statistical analysis, Elena Kost for her bioinformatical support, Dr. Daniel Wasner for his insights into visualization in R, Serena Richelle for the spinach and radish graphics, Sorya Lagnaux for the plant N uptake analysis, Prof. Dr. Daniel Buckley for hosting a research exchange during this project and for sharing his expertise on soil microbial ecology, Prof. Dr. Kai Udert for the Aurin fertilizer used in this study and Dr. Nikolas Hagemann for help in the greenhouse trial and providing the soil characterization. Dr. Felix Wettstein, Dr. Christa McArdell, Dr. Andreas Mccagnan, Dr. Andrea Rösch and Dr. Judith Riedo are kindly acknowledged for their insight on experimental set up and validation of the antibiotic extraction method.

## Author contributions

**Sarah van den Broek**: Conceptualization, Methodology, Formal analysis, Investigation, Data Curation, Writing – Original draft, Writing – Reviewing and Editing, Visualization. **Inna Nybom**: Conceptualization, Methodology, Formal analysis, Data Curation, Writing – Original draft, Writing – Reviewing and Editing, Supervision. **Rafaela Feola Conz**: Conceptualization, Methodology, Validation, Investigation, Writing - Reviewing and Editing, Supervision. **Yifei Sun**: Methodology, Writing – Reviewing and Editing. **Thomas Bucheli**: Resources, Validation, Writing – Reviewing and Editing. **Sebastian Doetterl**: Resources, Supervision, Writing – Reviewing and Editing. **Martin Hartmann**: Conceptualization, Methodology, Software, Validation, Resources, Writing – Reviewing and Editing, Supervision. **Gina Garland**: Conceptualization, Resources, Writing – Reviewing and Editing, Supervision, Project administration, Funding Acquisition.

## Data availability

The raw metabarcoding sequences were deposited in the European Nucleotide Archive (ENA) under accession number PRJEB95045. The data used to create Figure 5 is shared on Figshare doi:10.6084/m9.figshare.29820431.

## References

Abarenkov, K., Nilsson, R.H., Larsson, K.H., Taylor, A.F.S., May, T.W., Frøslev, T.G., Pawlowska, J., Lindahl, B., Põldmaa, K., Truong, C., Vu, D., Hosoya, T., Niskanen, T., Piirmann, T., Ivanov, F., Zirk, A., Peterson, M., Cheeke, T.E., Ishigami, Y., Jansson, A.T., Jeppesen, T.S., Kristiansson, E., Mikryukov, V., Miller, J.T., Oono, R., Ossandon, F.J., Paupério, J., Saar, I., Schigel, D., Suija, A., Tedersoo, L., Kõljalg, U., 2024. The UNITE database for molecular identification and taxonomic communication of fungi and other eukaryotes: sequences, taxa and classifications reconsidered. Nucleic Acids Res. 52, D791–D797. 10.1093/nar/gkad1039

Adobe Inc., 2019. Adobe Illustrator.

Ajibade, F.O., Yin, W.X., Guadie, A., Ajibade, T.F., Liu, Y., Kumwimba, M.N., Liu, W.Z., Han, J.L., Wang, H.C., Wang, A.J., 2023. Impact of biochar amendment on antibiotic removal and ARGs accumulation in constructed wetlands for low C/N wastewater treatment. Chem. Eng. J. 459, 141541. 10.1016/j.cej.2023.141541

Alcock, B.P., Raphenya, A.R., Lau, T.T.Y., Tsang, K.K., Bouchard, M., Edalatmand, A., Huynh, W., Nguyen, A.L. V., Cheng, A.A., Liu, S., Min, S.Y., Miroshnichenko, A., Tran, H.K., Werfalli, R.E., Nasir, J.A., Oloni, M., Speicher, D.J., Florescu, A., Singh, B., Faltyn, M., Hernandez-Koutoucheva, A., Sharma, A.N., Bordeleau, E., Pawlowski, A.C., Zubyk, H.L., Dooley, D., Griffiths, E., Maguire, F., Winsor, G.L., Beiko, R.G., Brinkman, F.S.L., Hsiao, W.W.L., Domselaar, G. V., McArthur, A.G., 2020. CARD 2020: Antibiotic resistome surveillance with the comprehensive antibiotic resistance database. Nucleic Acids Res. 48, D517–D525. 10.1093/nar/gkz935

Andrews, S., 2010. FastQC: A Quality Control tool for High Throughput Sequence Data.

Arseneault, T., Filion, M., 2017. Biocontrol through antibiosis: exploring the role played by subinhibitory concentrations of antibiotics in soil and their impact on plant pathogens. Can. J. Plant Pathol. 39, 267–274. 10.1080/07060661.2017.1354335

Avisar, D., Primor, O., Gozlan, I., Mamane, H., 2010. Sorption of sulfonamides and tetracyclines to montmorillonite clay. Water. Air. Soil Pollut. 209, 439–450. 10.1007/s11270-009-0212-8

Bartlett, A., Padfield, D., Lear, L., Bendall, R., Vos, M., 2022. A comprehensive list of bacterial pathogens infecting humans. Microbiol. (United Kingdom) 168, 1–8. 10.1099/mic.0.001269

Bengtsson-Palme, J., Hartmann, M., Eriksson, K.M., Pal, C., Thorell, K., Larsson, D.G.J., Nilsson, R.H., 2015. metaxa2: Improved identification and taxonomic classification of small and large subunit rRNA in metagenomic data. Mol. Ecol. Resour. 15, 1403–1414. 10.1111/1755-0998.12399

Bengtsson-Palme, J., Ryberg, M., Hartmann, M., Branco, S., Wang, Z., Godhe, A., De Wit, P., Sánchez-García, M., Ebersberger, I., de Sousa, F., Amend, A., Jumpponen, A., Unterseher, M., Kristiansson, E., Abarenkov, K., Bertrand, Y.J.K., Sanli, K., Eriksson, K.M., Vik, U., Veldre, V., Nilsson, R.H., 2013. Improved software detection and extraction of ITS1 and ITS2 from ribosomal ITS sequences of fungi and other eukaryotes for analysis of environmental sequencing data. Methods Ecol. Evol. 4, 914–919. 10.1111/2041-210X.12073

Bischel, H.N., Özel Duygan, B.D., Strande, L., McArdell, C.S., Udert, K.M., Kohn, T., 2015. Pathogens and pharmaceuticals in source-separated urine in eThekwini, South Africa. Water Res. 85, 57–65. 10.1016/j.watres.2015.08.022

Boss, B.L., Wanees, A.E., Zaslow, S.J., Normile, T.G., Izquierdo, J.A., 2022. Comparative genomics of the plant-growth promoting bacterium Sphingobium sp. strain AEW4 isolated from the rhizosphere of the beachgrass Ammophila breviligulata. BMC Genomics 23, 1–14. 10.1186/s12864-022-08738-8

Boxall, A.B.A., Johnson, P., Smith, E.J., Sinclair, C.J., Stutt, E., Levy, L.S., 2006. Uptake of veterinary medicines from soils into plants. J. Agric. Food Chem. 54, 2288–2297. 10.1021/jf053041t

Brown, L.P., Murray, R., Scott, A., Tien, Y.C., Lau, C.H.F., Tai, V., Topp, E., 2022. Responses of the Soil Bacterial Community, Resistome, and Mobilome to a Decade of Annual Exposure to Macrolide Antibiotics. Appl. Environ. Microbiol. 88. 10.1128/aem.00316-22

Brunhoferova, H., Venditti, S., Laczny, C.C., Lebrun, L., Hansen, J., 2022. Bioremediation of 27 Micropollutants by Symbiotic Microorganisms of Wetland Macrophytes. Sustain. 14. 10.3390/su14073944

Buffie, C.G., Pamer, E.G., 2013. Microbiota-mediated colonization resistance against intestinal pathogens. Nat. Rev. Immunol. 13, 790–801. 10.1038/nri3535

Bull, C.T., Boer, S.H. De, Denny, T.P., Firrao, G., Saux, M.F., Saddler, G.S., Scortichini, M., Stead, D.E., Takikawa, Y., 2010. Comprehensive List of Names of Plant Pathogenic Bacteria, 1980-2007. J. Plant Pathol. 92, 551–592.

Bünemann, E.K., Reimer, M., Smolders, E., Smith, S.R., Bigalke, M., Palmqvist, A., Brandt, K.K., Möller, K., Harder, R., Hermann, L., Speiser, B., Oudshoorn, F., Løes, A.K., Magid, J., 2024. Do contaminants compromise the use of recycled nutrients in organic agriculture? A review and synthesis of current knowledge on contaminant concentrations, fate in the environment and risk assessment. Sci. Total Environ. 912, 1–18. 10.1016/j.scitotenv.2023.168901

Burch, T.R., Sadowsky, M.J., LaPara, T.M., 2017. Effect of Different Treatment Technologies on the Fate of Antibiotic Resistance Genes and Class 1 Integrons when Residual Municipal Wastewater Solids are Applied to Soil. Environ. Sci. Technol. 51, 14225–14232. 10.1021/acs.est.7b04760

Carballo, M., Rodríguez, A., de la Torre, A., 2022. Phytotoxic Effects of Antibiotics on Terrestrial Crop Plants and Wild Plants: A Systematic Review. Arch. Environ. Contam. Toxicol. 82, 48–61. 10.1007/s00244-021-00893-5

Charif, D., Lobry, J.R., 2004. seqinr: Biological Sequences Retrieval and Analysis. CRAN Contrib. Packag. 10.32614/CRAN.PACKAGE.SEQINR

Chen, C., Li, J., Chen, P., Ding, R., Zhang, P., Li, X., 2014. Occurrence of antibiotics and antibiotic resistances in soils from wastewater irrigation areas in Beijing and Tianjin, China. Environ. Pollut. 193, 94–101. 10.1016/j.envpol.2014.06.005

Chen, J., Zhu, B., Zhang, Y., 2023. A Meta-Analysis on the Responses of Soil Microbial Biomass and Community Structure to Antibiotics. Appl. Soil Ecol. 184, 104786. 10.2139/ssrn.4224302

Chen, S., Zhou, Y., Chen, Y., Gu, J., 2018. Fastp: An ultra-fast all-in-one FASTQ preprocessor. Bioinformatics 34, i884–i890. 10.1093/bioinformatics/bty560

Chung, H.S., Lee, Y.J., Rahman, M.M., Abd El-Aty, A.M., Lee, H.S., Kabir, M.H., Kim, S.W., Park, B.J., Kim, J.E., Hacımüftüoğlu, F., Nahar, N., Shin, H.C., Shim, J.H., 2017. Uptake of the veterinary antibiotics chlortetracycline, enrofloxacin, and sulphathiazole from soil by radish. Sci. Total Environ. 605–606, 322–331. 10.1016/j.scitotenv.2017.06.231

Cleary, D.W., Bishop, A.H., Zhang, L., Topp, E., Wellington, E.M.H., Gaze, W.H., 2016. Long-term antibiotic exposure in soil is associated with changes in microbial community structure and prevalence of class 1 integrons. FEMS Microbiol. Ecol. 92, 1–7. 10.1093/femsec/fiw159

Cui, M., Yu, S., Yu, Y., Chen, X., Li, J., 2022. Responses of cherry radish to different types of microplastics in the presence of oxytetracycline. Plant Physiol. Biochem. 191, 1–9. 10.1016/j.plaphy.2022.09.012

Cycoń, M., Mrozik, A., Piotrowska-Seget, Z., 2019. Antibiotics in the soil environment—degradation and their impact on microbial activity and diversity. Front. Microbiol. 10, 1–45. 10.3389/fmicb.2019.00338

de los Santos, E., Laviña, M., Poey, M.E., 2021. Strict relationship between class 1 integrons and resistance to sulfamethoxazole in Escherichia coli. Microb. Pathog. 161. 10.1016/j.micpath.2021.105206

Dean, A., Morris, M., Stufken, J., Bingham, D., 2015. Handbook of Design and Analysis of Experiments, Handbook of Design and Analysis of Experiments. 10.1201/b18619

Delgado-Baquerizo, M., Hu, H.W., Maestre, F.T., Guerra, C.A., Eisenhauer, N., Eldridge, D.J., Zhu, Y.G., Chen, Q.L., Trivedi, P., Du, S., Makhalanyane, T.P., Verma, J.P., Gozalo, B., Ochoa, V., Asensio, S., Wang, L., Zaady, E., Illán, J.G., Siebe, C., Grebenc, T., Zhou, X., Liu, Y.R., Bamigboye, A.R., Blanco-Pastor, J.L., Duran, J., Rodríguez, A., Mamet, S., Alfaro, F., Abades, S., Teixido, A.L., Peñaloza-Bojacá, G.F., Molina-Montenegro, M.A., Torres-Díaz, C., Perez, C., Gallardo, A., García-Velázquez, L., Hayes, P.E., Neuhauser, S., He, J.Z., 2022. The global distribution and environmental drivers of the soil antibiotic resistome. Microbiome 10, 1–15. 10.1186/s40168-022-01405-w

Delgado-Baquerizo, M., Maestre, F.T., Reich, P.B., Jeffries, T.C., Gaitan, J.J., Encinar, D., Berdugo, M., Campbell, C.D., Singh, B.K., 2016. Microbial diversity drives multifunctionality in terrestrial ecosystems. Nat. Commun. 7, 1–8. 10.1038/ncomms10541

Demoling, L.A., Bååth, E., Greve, G., Wouterse, M., Schmitt, H., 2009. Effects of sulfamethoxazole on soil microbial communities after adding substrate. Soil Biol. Biochem. 41, 840–848. 10.1016/j.soilbio.2009.02.001

Ding, G.C., Radl, V., Schloter-Hai, B., Jechalke, S., Heuer, H., Smalla, K., Schloter, M., 2014. Dynamics of soil bacterial communities in response to repeated application of manure containing sulfadiazine. PLoS One 9. 10.1371/journal.pone.0092958

Edgar, R., 2016. SINTAX: a simple non-Bayesian taxonomy classifier for 16S and ITS sequences. bioRxiv 074161.

Edgar, R.C., 2016a. UNOISE2: improved error-correction for Illumina 16S and ITS amplicon sequencing. bioRxiv 081257.

Edgar, R.C., 2016b. UCHIME2: improved chimera prediction for amplicon sequencing. bioRxiv 074252.

Fang, H., Han, L., Cui, Y., Xue, Y., Cai, L., Yu, Y., 2016. Changes in soil microbial community structure and function associated with degradation and resistance of carbendazim and chlortetracycline during repeated treatments. Sci. Total Environ. 572, 1203–1212. 10.1016/j.scitotenv.2016.08.038

Fiaz, M., Ahmed, I., Hassan, S.M.U., Niazi, A.K., Khokhar, M.F., Zeshan, Farooq, M.A., Arshad, M., 2023. Antibiotics induced changes in nitrogen metabolism and antioxidative enzymes in mung bean (Vigna radiata). Sci. Total Environ. 873, 162449. 10.1016/j.scitotenv.2023.162449

Frey, B., Rime, T., Phillips, M., Stierli, B., Hajdas, I., Widmer, F., Hartmann, M., 2016. Microbial diversity in European alpine permafrost and active layers. FEMS Microbiol. Ecol. 92, 1–17. 10.1093/femsec/fiw018

Frey, L., Tanunchai, B., Glaser, B., 2022. Antibiotics residues in pig slurry and manure and its environmental contamination potential. A meta-analysis. Agron. Sustain. Dev. 42. 10.1007/s13593-022-00762-y

Gao, Q., Gao, S., Bates, C., Zeng, Y., Lei, J., Su, H., Dong, Q., Qin, Z., Zhao, J., Zhang, Q., Ning, D., Huang, Y., Zhou, J., Yang, Y., 2021. The microbial network property as a bio-indicator of antibiotic transmission in the environment. Sci. Total Environ. 758, 143712. 10.1016/j.scitotenv.2020.143712

Garbini, G.L., Grenni, P., Rauseo, J., Patrolecco, L., Pescatore, T., Spataro, F., Barra Caracciolo, A., 2022. Insights into structure and functioning of a soil microbial community amended with cattle manure digestate and sulfamethoxazole. J. Soils Sediments 22, 2158–2173. 10.1007/s11368-022-03222-y

Gattinger, D., Schlenz, V., Weil, T., Sattler, B., 2024. From remote to urbanized: Dispersal of antibiotic-resistant bacteria under the aspect of anthropogenic influence. Sci. Total Environ. 924, 171532. 10.1016/j.scitotenv.2024.171532

Graves, S., Piepho, H.-P., Selzer, L., 2024. multcompView: Visualizations of Paired Comparisons.

Halawa, E.M., Fadel, M., Al-Rabia, M.W., Behairy, A., Nouh, N.A., Abdo, M., Olga, R., Fericean, L., Atwa, A.M., El-Nablaway, M., Abdeen, A., 2023. Antibiotic action and resistance: updated review of mechanisms, spread, influencing factors, and alternative approaches for combating resistance. Front. Pharmacol. 14, 1–17. 10.3389/fphar.2023.1305294

Hansson, K., Sundström, L., Pelletier, A., Roy, P.H., 2002. IntI2 integron integrase in Tn7. J. Bacteriol. 184, 1712–1721. 10.1128/JB.184.6.1712-1721.2002

Harrell, J.F., 2025. Hmisc: Harrell Miscellaneous.

Hothorn, T., Zeileis, A., Farebrother, R.W., Cummins, C., 2022. Testing Linear Regression Models [R package lmtest version 0.9-40]. CRAN Contrib. Packag. 10.32614/CRAN.PACKAGE.LMTEST

Hou, Q., Kolodkin-Gal, I., 2020. Harvesting the complex pathways of antibiotic production and resistance of soil bacilli for optimizing plant microbiome. FEMS Microbiol. Ecol. 96, 1–12. 10.1093/femsec/fiaa142

Hu, H.W., Wang, J.T., Li, J., Li, J.J., Ma, Y.B., Chen, D., He, J.Z., 2016. Field-based evidence for copper contamination induced changes of antibiotic resistance in agricultural soils. Environ. Microbiol. 18, 3896–3909. 10.1111/1462-2920.13370

Jia, W.L., Gao, F.Z., Song, C., Chen, C.E., Ma, C.X., White, J.C., Ying, G.G., 2024. Swine wastewater co-exposed with veterinary antibiotics enhanced the antibiotic resistance of endophytes in radish (Raphanus sativus L.). Environ. Pollut. 362, 125040. 10.1016/j.envpol.2024.125040

Jiao, S., Chen, W., Wei, G., 2019. Resilience and assemblage of soil microbiome in response to chemical contamination combined with plant growth. Appl. Environ. Microbiol. 85. 10.1128/AEM.02523-18

John, F., Weisberg, S., Price, B., Adler, D., Bates, D., Baud-bovy, G., Bolker, B., Ellison, S., Graves, S., Krivitsky, P., Laboissiere, R., Maechler, M., Monette, G., Murdoch, D., Ogle, D., Ripley, B., Venables, W., Walker, S., Winsemius, D., 2020. Package ‘car.’

Keenum, I., Liguori, K., Calarco, J., Davis, B.C., Milligan, E., Harwood, V.J., Pruden, A., 2022. A framework for standardized qPCR-targets and protocols for quantifying antibiotic resistance in surface water, recycled water and wastewater. Crit. Rev. Environ. Sci. Technol. 52, 4395–4419. 10.1080/10643389.2021.2024739

Kerrn, M.B., Klemmensen, T., Frimodt-Möller, N., Espersen, F., 2002. Susceptibility of Danish Escherichia coli strains isolated from urinary tract infections and bacteraemia, and distribution of sul genes conferring sulphonamide resistance. J. Antimicrob. Chemother. 50, 513–516. 10.1093/jac/dkf164

Khan, N.M., Imran, M., Ashraf, M., Arshad, H., Awan, A.R., 2022. Oxytetracycline and Ciprofloxacin Antibiotics Exhibit Contrasting Effects on Soil Microflora, Nitrogen Uptake, Growth, and Yield of Wheat (Triticum aestivum L.). J. Soil Sci. Plant Nutr. 22, 3788–3797. 10.1007/s42729-022-00927-4

Kodešová, R., Chroňáková, A., Grabicová, K., Kočárek, M., Schmidtová, Z., Frková, Z., Vojs Staňová, A., Nikodem, A., Klement, A., Fér, M., Grabic, R., 2020. How microbial community composition, sorption and simultaneous application of six pharmaceuticals affect their dissipation in soils. Sci. Total Environ. 746. 10.1016/j.scitotenv.2020.141134

Krupka, M., Piotrowicz-Cieślak, A.I., Michalczyk, D.J., 2022. Effects of antibiotics on the photosynthetic apparatus of plants. J. Plant Interact. 17, 96–104. 10.1080/17429145.2021.2014579

Kümmerer, K., 2009. Antibiotics in the aquatic environment - A review - Part I. Chemosphere 75, 417–434. 10.1016/j.chemosphere.2008.11.086

Langmead, B., Salzberg, S.L., 2012. Fast gapped-read alignment with Bowtie 2. Nat. Methods 9, 357–359. 10.1038/nmeth.1923

Larsson, D.G.J., Flach, C.F., 2022. Antibiotic resistance in the environment. Nat. Rev. Microbiol. 20, 257–269. 10.1038/s41579-021-00649-x

Lau, C.H.F., Tien, Y.C., Stedtfeld, R.D., Topp, E., 2020. Impacts of multi-year field exposure of agricultural soil to macrolide antibiotics on the abundance of antibiotic resistance genes and selected mobile genetic elements. Sci. Total Environ. 727, 138520. 10.1016/j.scitotenv.2020.138520

Laureti, L., Matic, I., Gutierrez, A., 2013. Bacterial responses and genome instability induced by subinhibitory concentrations of antibiotics. Antibiotics 2, 100–114. 10.3390/antibiotics2010100

Lei, H., Zhang, J., Huang, J., Shen, D., Li, Y., Jiao, R., Zhao, R., Li, X., Lin, L., Li, B., 2023. New insights into lincomycin biodegradation by Conexibacter sp. LD01: Genomics characterization, biodegradation kinetics and pathways. J. Hazard. Mater. 441, 129824. 10.1016/j.jhazmat.2022.129824

Li, L., Li, T., Liu, Y., Li, Lina, Huang, X., Xie, J., 2023. Effects of antibiotics stress on root development, seedling growth, antioxidant status and abscisic acid level in wheat (Triticum aestivum L.). Ecotoxicol. Environ. Saf. 252. 10.1016/j.ecoenv.2023.114621

Li, P., Tedersoo, L., Crowther, T.W., Dumbrell, A.J., Dini-Andreote, F., Bahram, M., Kuang, L., Li, T., Wu, M., Jiang, Y., Luan, L., Saleem, M., de Vries, F.T., Li, Z., Wang, B., Jiang, J., 2023. Fossil-fuel-dependent scenarios could lead to a significant decline of global plant-beneficial bacteria abundance in soils by 2100. Nat. Food 4, 996–1006. 10.1038/s43016-023-00869-9

Li, T., Cao, X., Wu, Z., Liu, J., Hu, B., Chen, H., Li, B., 2023. Biotransformation of nitrogen and tetracycline by counter-diffusion biofilm system: Multiple metabolic pathways, mechanism, and slower resistance genes enrichment. Chem. Eng. J. 474, 145637. 10.1016/j.cej.2023.145637

Li, X., Lu, Z., Wu, B., Xie, H., Liu, G., 2024. Antibiotics and antibiotic resistance genes removal in biological aerated filter. Bioresour. Technol. 395, 130392. 10.1016/j.biortech.2024.130392

Liao, H., Lu, X., Rensing, C., Friman, V.P., Geisen, S., Chen, Z., Yu, Z., Wei, Z., Zhou, S., Zhu, Y., 2018. Hyperthermophilic Composting Accelerates the Removal of Antibiotic Resistance Genes and Mobile Genetic Elements in Sewage Sludge. Environ. Sci. Technol. 52, 266–276. 10.1021/acs.est.7b04483

Lin, K., Gan, J., 2011. Sorption and degradation of wastewater-associated non-steroidal anti-inflammatory drugs and antibiotics in soils. Chemosphere 83, 240–246. 10.1016/j.chemosphere.2010.12.083

Liu, F., Ying, G.G., Tao, R., Zhao, J.L., Yang, J.F., Zhao, L.F., 2009a. Effects of six selected antibiotics on plant growth and soil microbial and enzymatic activities. Environ. Pollut. 157, 1636–1642. 10.1016/j.envpol.2008.12.021

Liu, F., Ying, G.G., Tao, R., Zhao, J.L., Yang, J.F., Zhao, L.F., 2009b. Effects of six selected antibiotics on plant growth and soil microbial and enzymatic activities. Environ. Pollut. 157, 1636–1642. 10.1016/j.envpol.2008.12.021

Liu, W., Li, B., Chu, H., Zhang, Z., Luo, L., Ma, W., Yang, S., Guo, Q., 2017. Rapid detection of mutations in erm(41) and rrl associated with clarithromycin resistance in Mycobacterium abscessus complex by denaturing gradient gel electrophoresis. J. Microbiol. Methods 143, 87–93. 10.1016/j.mimet.2017.10.010

Longepierre, M., Widmer, F., Keller, T., Weisskopf, P., Colombi, T., Six, J., Hartmann, M., 2021. Limited resilience of the soil microbiome to mechanical compaction within four growing seasons of agricultural management. ISME Commun. 1, 1–13. 10.1038/s43705-021-00046-8

Luo, G., Li, L., Friman, V.P., Guo, J., Guo, S., Shen, Q., Ling, N., 2018. Organic amendments increase crop yields by improving microbe-mediated soil functioning of agroecosystems: A meta-analysis. Soil Biol. Biochem. 124, 105–115. 10.1016/j.soilbio.2018.06.002

Luo, Y., Mao, D., Rysz, M., Zhou, Q., Zhang, H., Xu, L., Alvarez, P.J.J., 2010. Trends in antibiotic resistance genes occurrence in the Haihe River, China. Environ. Sci. Technol. 44, 7220–7225. 10.1021/es100233w

Mahdi, I., Fahsi, N., Hijri, M., Sobeh, M., 2022. Antibiotic resistance in plant growth promoting bacteria: A comprehensive review and future perspectives to mitigate potential gene invasion risks. Front. Microbiol. 13, 1–22. 10.3389/fmicb.2022.999988

Marti, E., Balcázar, J.L., 2013. Real-time PCR assays for quantification of qnr genes in environmental water samples and chicken feces. Appl. Environ. Microbiol. 79, 1743–1745. 10.1128/AEM.03409-12

Martin, M., 2011. Cutadapt removes adapter sequences from high-throughput sequencing reads. EMBnet J 17, 10–12. 10.14806/ej.17.1.200

Martinez Arbizu, P., 2020. pairwiseAdonis: Pairwise multilevel comparison using adonis.

McFarland, J.W., Berger, C.M., Froshauer, S.A., Hayashi, S.F., Hecker, S.J., Jaynes, B.H., Jefson, M.R., Kamicker, B.J., Lipinski, C.A., Lundy, K.M., Reese, C.P., Vu, C.B., 1997. Quantitative structure-activity relationships among macrolide antibacterial agents: In vitro and in vivo potency against Pasteurella multocida. J. Med. Chem. 40, 1340–1346. 10.1021/jm960436i

Minden, V., Deloy, A., Volkert, A.M., Leonhardt, S.D., Pufal, G., 2017. Antibiotics impact plant traits, even at small concentrations. AoB Plants 9. 10.1093/aobpla/plx010

Mohy-u-Din, N., Farhan, M., Wahid, A., Ciric, L., Sharif, F., 2023. Human health risk estimation of antibiotics transferred from wastewater and soil to crops. Environ. Sci. Pollut. Res. 30, 20601–20614. 10.1007/s11356-022-23412-y

Nafees, M., Qiu, L., Alomrani, S.O., Ahmad, Z., Yin, Y., Sallah A, A.H., Alshehri, M.A., Ali, S., Guo, H., 2025. Enhancing spinach growth and soil microbial health under sulfadiazine and polypropylene exposure through zinc fortification. Environ. Technol. Innov. 38, 104186. 10.1016/j.eti.2025.104186

National Center for Biotechnology Information, 2004. Gene [Internet]. Bethesda (MD): National Library of Medicine (US), National Center for Biotechnology Information [WWW Document]. URL https://www.ncbi.nlm.nih.gov/

Neuwirth, E., 2011. Package “RColorBrewer.” Phys. Rev. D - Part. Fields, Gravit. Cosmol. 84. 10.1103/PhysRevD.84.026007

Nguyen, N.H., Song, Z., Bates, S.T., Branco, S., Tedersoo, L., Menke, J., Schilling, J.S., Kennedy, P.G., 2016. FUNGuild: An open annotation tool for parsing fungal community datasets by ecological guild. Fungal Ecol. 20, 241–248. 10.1016/j.funeco.2015.06.006

Nowara, A., Burhenne, J., Spiteller, M., 1997. Binding of Fluoroquinolone Carboxylic Acid Derivatives to Clay Minerals. J. Agric. Food Chem. 45, 1459–1463. 10.1021/jf960215l

Oksanen, A.J., Blanchet, F.G., Friendly, M., Kindt, R., Legendre, P., Mcglinn, D., Minchin, P.R., Hara, R.B.O., Simpson, G.L., Solymos, P., Stevens, M.H.H., Szoecs, E., 2012. Vegan: Community Ecology Package.

Osose, E.B., Okafor, C.O.O., Nnenna, O.E., Ndidi, O.K., Ikechukwu, O., Francisca, A.U., Felicia, O.N., Chinyere, N.A., Romanus, I.I., 2025. Multidrug resistant Salmonella species harboring sulfonamide (sul1 and sul2) resistant gene isolated from typhoid patients in a university teaching hospital – A threat to public health. Total Environ. Microbiol. 1, 100021. 10.1016/j.temicr.2025.100021

Pan, M., Chu, L.M., 2016. Phytotoxicity of veterinary antibiotics to seed germination and root elongation of crops. Ecotoxicol. Environ. Saf. 126, 228–237. 10.1016/j.ecoenv.2015.12.027

Pedersen, T.L., 2025. patchwork: The Composer of Plots.

Pruesse, E., Quast, C., Knittel, K., Fuchs, B.M., Ludwig, W., Peplies, J., Glöckner, F.O., 2007. SILVA: A comprehensive online resource for quality checked and aligned ribosomal RNA sequence data compatible with ARB. Nucleic Acids Res. 35, 7188–7196. 10.1093/nar/gkm864

Qin, S., Xiao, W., Zhou, C., Pu, Q., Deng, X., Lan, L., Liang, H., Song, X., Wu, M., 2022. Pseudomonas aeruginosa: pathogenesis, virulence factors, antibiotic resistance, interaction with host, technology advances and emerging therapeutics. Signal Transduct. Target. Ther. 7, 1–27. 10.1038/s41392-022-01056-1

R Core Team, 2021. A Language and Environment for Statistical Computing.

Ransirini, A.M., Elżbieta, M.S., Joanna, G., Bartosz, K., Wojciech, T., Agnieszka, B., Magdalena, U., 2024. Fertilizing drug resistance: Dissemination of antibiotic resistance genes in soil and plant bacteria under bovine and swine slurry fertilization. Sci. Total Environ. 946. 10.1016/j.scitotenv.2024.174476

Ren, J., Lu, H., Lu, S., Huang, Z., 2024. Impacts of sulfamethoxazole stress on vegetable growth and rhizosphere bacteria and the corresponding mitigation mechanism. Front. Bioeng. Biotechnol. 12, 1–11. 10.3389/fbioe.2024.1303670

Rice, L.B., 2008. Federal funding for the study of antimicrobial resistance in nosocomial pathogens: No ESKAPE. J. Infect. Dis. 197, 1079–1081. 10.1086/533452

Richter, L., du Plessis, E.M., Duvenage, S., Korsten, L., 2020. Occurrence, Phenotypic and Molecular Characterization of Extended-Spectrum- and AmpC-β-Lactamase Producing Enterobacteriaceae Isolated From Selected Commercial Spinach Supply Chains in South Africa. Front. Microbiol. 11, 1–10. 10.3389/fmicb.2020.00638

Rognes, T., Flouri, T., Nichols, B., Quince, C., Mahé, F., 2016. VSEARCH: A versatile open source tool for metagenomics. PeerJ 2016, 1–22. 10.7717/peerj.2584

Romero, F., Hilfiker, S., Edlinger, A., Held, A., Hartman, K., Labouyrie, M., van der Heijden, M.G.A., 2023. Soil microbial biodiversity promotes crop productivity and agro-ecosystem functioning in experimental microcosms. Sci. Total Environ. 885. 10.1016/j.scitotenv.2023.163683

Rosewarne, C.P., Pettigrove, V., Stokes, H.W., Parsons, Y.M., 2010. Class 1 integrons in benthic bacterial communities: Abundance, association with Tn402-like transposition modules and evidence for coselection with heavy-metal resistance. FEMS Microbiol. Ecol. 72, 35–46. 10.1111/j.1574-6941.2009.00823.x

RStudio Team, 2020. RStudio: Integrated Development for R.

Sah, S., Krishnani, S., Singh, R., 2021. Pseudomonas mediated nutritional and growth promotional activities for sustainable food security. Curr. Res. Microb. Sci. 2, 100084. 10.1016/j.crmicr.2021.100084

Sarmah, A.K., Meyer, M.T., Boxall, A.B.A., 2006. A global perspective on the use, sales, exposure pathways, occurrence, fate and effects of veterinary antibiotics (VAs) in the environment. Chemosphere 65, 725–759. 10.1016/j.chemosphere.2006.03.026

Schloss, P.D., 2024. Rarefaction is currently the best approach to control for uneven sequencing effort in amplicon sequence analyses. mSphere 9, 1–20. 10.1128/msphere.00354-23

Schloss, P.D., 2023. Waste not, want not: revisiting the analysis that called into question the practice of rarefaction. mSphere 0, 1–22. 10.1128/msphere.00355-23

Shen, Y., Ryser, E.T., Li, H., Zhang, W., 2021. Bacterial community assembly and antibiotic resistance genes in the lettuce-soil system upon antibiotic exposure. Sci. Total Environ. 778, 146255. 10.1016/j.scitotenv.2021.146255

Shi, X., Zhang, S., Zhang, Y., Geng, Y., Wang, L., Peng, Y., He, Z., 2022. Novel and simple analytical method for simultaneous determination of sulfonamide, quinolone, tetracycline, macrolide, and chloramphenicol antibiotics in soil. Anal. Bioanal. Chem. 414, 6497–6506. 10.1007/s00216-022-04206-0

Sidhu, H., O’Connor, G., Kruse, J., 2019. Plant toxicity and accumulation of biosolids-borne ciprofloxacin and azithromycin. Sci. Total Environ. 648, 1219–1226. 10.1016/j.scitotenv.2018.08.218

Stephens, R., 1956. Acidity Antibiotics 3, 12–15.

Stoob, K., Singer, H.P., Mueller, S.R., Schwarzenbach, R.P., Stamm, C.H., 2007. Dissipation and transport of veterinary sulfonamide antibiotics after manure application to grassland in a small catchment. Environ. Sci. Technol. 41, 7349–7355. 10.1021/es070840e

Tedersoo, L., Lindahl, B., 2016. Fungal identification biases in microbiome projects. Environ. Microbiol. Rep. 8, 774–779. 10.1111/1758-2229.12438

Thiele-Bruhn, S., 2003. Pharmaceutical antibiotic compounds in soils - A review. J. Plant Nutr. Soil Sci. 166, 145–167. 10.1002/jpln.200390023

Tian, L., Sun, H., Dong, X., Wang, J., Huang, Y., Sun, S., 2022. Effects of swine wastewater irrigation on soil properties and accumulation of heavy metals and antibiotics. J. Soils Sediments 22, 889–904. 10.1007/s11368-021-03106-7

Tian, S., You, L., Huang, X., Liu, C., Su, J.Q., 2024. Efficient sulfamethoxazole biotransformation and detoxification by newly isolated strain Hydrogenophaga sp. SNF1 via a ring ortho-hydroxylation pathway. J. Hazard. Mater. 480, 136113. 10.1016/j.jhazmat.2024.136113

Timmerer, U., Lehmann, L., Schnug, E., Bloem, E., 2020. Toxic effects of single antibiotics and antibiotics in combination on germination and growth of Sinapis alba L. Plants 9. 10.3390/plants9010107

Trimmer, J.T., Cusick, R.D., Guest, J.S., 2017. Amplifying Progress toward Multiple Development Goals through Resource Recovery from Sanitation. Environ. Sci. Technol. 51, 10765–10776. 10.1021/acs.est.7b02147

Urra, J., Alkorta, I., Garbisu, C., 2019. Potential benefits and risks for soil health derived from the use of organic amendments in agriculture. Agronomy 9, 1–23. 10.3390/agronomy9090542

van den Broek, S., Nybom, I., Hartmann, M., Doetterl, S., Garland, G., 2024. Opportunities and challenges of using human excreta-derived fertilizers in agriculture: A review of suitability, environmental impact and societal acceptance. Sci. Total Environ. 957, 177306. 10.1016/j.scitotenv.2024.177306

Van Elsas, J.D., Chiurazzi, M., Mallon, C.A., Elhottova, D., Krištůfek, V., Salles, J.F., 2012. Microbial diversity determines the invasion of soil by a bacterial pathogen. Proc. Natl. Acad. Sci. U. S. A. 109, 1159–1164. 10.1073/pnas.1109326109

Vives-Peris, V., de Ollas, C., Gómez-Cadenas, A., Pérez-Clemente, R.M., 2020. Root exudates: from plant to rhizosphere and beyond. Plant Cell Rep. 39, 3–17. 10.1007/s00299-019-02447-5

Waglechner, N., Wright, G.D., 2017. Antibiotic resistance: It’s bad, but why isn’t it worse? BMC Biol. 15, 1–8. 10.1186/s12915-017-0423-1

Wash, P., Batool, A., Mulk, S., Nazir, S., Yasmin, H., Mumtaz, S., Alyemeni, M.N., Kaushik, P., Hassan, M.N., 2022. Prevalence of Antimicrobial Resistance and Respective Genes among Bacillus spp., a Versatile Bio-Fungicide. Int. J. Environ. Res. Public Health 19. 10.3390/ijerph192214997

Wei, T., Simko, V., 2024. R package “corrplot”: Visualization of a Correlation Matrix.

Wickham, H., Chang, W., 2016. Create elegant data visualisations using the grammar of graphics.

Wu, J., Wang, J., Li, Z., Guo, S., Li, K., Xu, P., Ok, Y.S., Jones, D.L., Zou, J., 2023. Antibiotics and antibiotic resistance genes in agricultural soils: A systematic analysis. Crit. Rev. Environ. Sci. Technol. 53, 847–864. 10.1080/10643389.2022.2094693

Xie, W.Y., McGrath, S.P., Su, J.Q., Hirsch, P.R., Clark, I.M., Shen, Q., Zhu, Y.G., Zhao, F.J., 2016. Long-term impact of field applications of sewage sludge on soil antibiotic resistome. Environ. Sci. Technol. 50, 12602–12611. 10.1021/acs.est.6b02138

Yang, J.F., Ying, G.G., Zhou, L.J., Liu, S., Zhao, J.L., 2009. Dissipation of oxytetracycline in soils under different redox conditions. Environ. Pollut. 157, 2704–2709. 10.1016/j.envpol.2009.04.031

Yang, Q., Gao, Y., Ke, J., Show, P.L., Ge, Y., Liu, Y., Guo, R., Chen, J., 2021. Antibiotics: An overview on the environmental occurrence, toxicity, degradation, and removal methods. Bioengineered 12, 7376–7416. 10.1080/21655979.2021.1974657

Yin, L., Wang, X., Li, Y., Liu, Z., Mei, Q., Chen, Z., 2023. Uptake of the Plant Agriculture-Used Antibiotics Oxytetracycline and Streptomycin by Cherry Radish─Effect on Plant Microbiome and the Potential Health Risk. J. Agric. Food Chem. 71, 4561–4570. 10.1021/acs.jafc.3c01052

Zalewska, M., Błażejewska, A., Czapko, A., Popowska, M., 2021. Antibiotics and Antibiotic Resistance Genes in Animal Manure – Consequences of Its Application in Agriculture. Front. Microbiol. 12. 10.3389/fmicb.2021.610656

Zhang, Y., Zheng, X., Xu, X., Cao, L., Zhang, Haiyun, Zhang, Hanlin, Li, S., Zhang, J., Bai, N., Lv, W., Cao, X., 2022. Straw return promoted the simultaneous elimination of sulfamethoxazole and related antibiotic resistance genes in the paddy soil. Sci. Total Environ. 806, 150525. 10.1016/j.scitotenv.2021.150525

Zhou, S.Y.D., Wei, M.Y., Giles, M., Neilson, R., Zheng, F., Zhang, Q., Zhu, Y.G., Yang, X.R., 2020. Prevalence of Antibiotic Resistome in Ready-to-Eat Salad. Front. Public Heal. 8, 1–9. 10.3389/fpubh.2020.00092

